# Antagonism between bacteriophages and macrophages decreases efficacy of a bacteriophage cocktail and increases bacteriophage resistance

**DOI:** 10.1101/2025.05.16.654446

**Authors:** Meaghan Castledine, Zuzanna Szczutkowska, Andrew Matthews, Sarah K Walsh, Rai Lewis, Suzanne Kay, Janet A Willment, Gordon D. Brown, Angus Buckling

## Abstract

Phage therapy, the use of viruses that infect bacteria (bacteriophages), is a promising complement to antibiotics during the antimicrobial resistance crisis, but treatment success is very variable. Evolution of bacterial resistance to bacteriophages and bacteriophage counter-resistance (coevolution) during therapy may explain some of this variation, the dynamics of which may be affected by interactions with the patient’s immune system. Here, we examine how a pathogenic bacterium, *Pseudomonas aeruginosa* coevolves with two clinically relevant bacteriophages (14-1 and PNM) when in the presence of macrophages (RAW 264.7 cell line). We show macrophages reduced the rate by which bacteria were killed by bacteriophages, likely by reducing bacteria-bacteriophage contact rates. Over evolutionary time-scales, macrophages increased the proportion of bacteriophage resistant bacteria compared to where macrophages were absent. These differences in resistance rates were likely driven by the early advantage in density offered by macrophages to bacteria, and exclusion of PNM from the bacteriophage cocktail which otherwise increased in frequency in the absence of macrophages. Consequently, macrophages significantly altered the short- and long-term efficacy of a bacteriophage cocktail. In line with a growing body of work, our results suggest that the patient’s immune system can reduce the efficacy of phage therapy, potentially driving variable outcomes in therapy success in patients.

**Significance statement:** Phage therapy, the use of viruses that infect bacteria (bacteriophages), is a promising complement to antibiotics during the antimicrobial resistance crisis. However, treatment success is very variable. One, often overlooked, variable is the immune system and how this influences bacteriophage efficacy, and how bacteria evolve resistance to bacteriophages. We find that macrophages reduce the rate by which bacteria are killed by bacteriophages. Resistance to bacteriophages also increased in the presence of macrophages, showing macrophages affect the short- and long-term efficacy of phage therapy. These results highlight the importance of the immune system in phage therapy, and the need for more research in this area.

## Introduction

In 2019 4.95 million deaths were associated with antimicrobial resistant (AMR) infections, with this figure expected to double by 2050 (1). To combat the AMR crisis, new treatments are required such as phage therapy (2). Phage therapy uses viruses (bacteriophages) that specifically infect and kill bacteria to treat bacterial infections with reported success in patients and animal models (2–4). As bacteriophages are typically species-specific and self-amplifying at infection sites, they can target pathogenic bacteria without disturbing microbiomes (5, 6). However, phage therapy outcomes are highly diverse and clinical trials have produced mixed results for determining efficacy in pathogen eradication (2, 7, 8). The immune system is likely to be responsible for much of this unpredictability with patients varying in immunocompromised states and pathogens varying in their ability to evade different immune components. Although the immune system is suggested to be key for phage therapy success (9, 10), little experimental work has been conducted to test how immune components interact with phage therapeutics. What work has been conducted has yielded mixed-results with immune components both aiding and hindering phage therapy success (3, 11, 12).

The effect of the immune system on phage therapy efficacy is further complicated by (co)evolution. Bacterial pathogens can rapidly evolve resistance to bacteriophages (13), which has been demonstrated *in vitro* (14) and in clinical case studies (7, 15), and bacteriophages can evolve to overcome resistance (14, 16). Recent work suggests that synergy with the immune system and bacteriophages is even more important for treatment success when (co)evolution occurs, with immune cells clearing resistant bacteria and viruses clearing susceptible populations (17, 18). Mice that can produce neutrophils had significantly higher survival probabilities than immuno-compromised mice when treated for an infection with phage therapy; although this was only mathematically linked to bacteriophage resistance (17). Bacteriophage resistance has been associated with reduced bacterial survival in insect haemolymph (19) and increased phagocytosis (20). However, bacteriophage resistance has also been found to positively covary with resistance to phagocytosis (11). These different outcomes may in part be explained by the types of resistance mutation (21). For example, biofilm production is a common mechanism of bacteriophage resistance (22, 23), that can also confer resistance to phagocytes, and increase release of pro-inflammatory cytokines (21). Additionally, bacteriophages can be internalised by phagocytes (24, 25) (or bound by antibodies (26)) and so synergy between the immune system and bacteriophages may depend on the relative rates of replication and phagocytosis between pathogens and viruses (3). The mixed evidence regarding whether immune cells and bacteriophages are synergistic and the limited research examining how bacteriophage resistance affects these dynamics warrants further research.

Immune cells may not only affect the evolution of bacterial resistance, but also how bacteriophages evolve to overcome resistance. While no study has examined bacteria-bacteriophage coevolution with immune cells, studies have examined how other antagonists influence these dynamics. Protists, for instance, can limit rates of bacteria-phage coevolution by preventing an accumulation of resistance and infectivity and instead shifting dynamics such that selection is on standing variation (not primarily new mutations) (27). Additionally, any antagonist that limits bacterial replication (e.g. plasmids, competitors) can reduce the probability of bacteriophage resistance as mutation rates decline (28, 29). However, whether these effects are applicable to immune cells’ influence on bacteria-phage coevolution is unstudied.

Here, we address this knowledge gap using macrophages (RAW 264.7) and assess their impact on the evolution of bacteriophage resistance and bacteria-phage (co)evolution. Macrophages are a generally resilient cell line and are one of the first cell types to respond during an infection (30), thereby making these cells an attractive and applicable model system for understanding immune function. Through tissue culturing *in vitro*, RAW 264.7 macrophages can be used experimentally in highly controlled conditions, allowing researchers to examine how immune cells may interact with pathogens under different experimental treatments. As such, we passage a common bacterial pathogen, *Pseudomonas aeruginosa* (mucoid strain P573), with bacteriophages 14-1 and PNM through flasks containing macrophages and examine how macrophages affect bacteria-bacteriophage coevolution and the resultant impact on the efficacy of bacteriophages in controlling bacterial densities. We further considered how bacteriophage resistance affected biofilm production, with P573 being a mucoid strain, and subsequently any changes in pro-inflammatory cytokines released by macrophages as a marker for the effects of resistance on immune function.

## Results

### Macrophages and bacteriophages interact antagonistically

Before examining bacteria and bacteriophage coevolutionary interactions, we first considered how the presence of macrophages impacts bacteria-phage ecological interactions. To this end, bacteria were cultured in the presence and absence of macrophages and bacteriophages for 8 hrs (before bacteria reach carrying capacity).

When bacteria were cultured with bacteriophages alone, bacteriophages decreased bacterial densities to below detectable levels in all replicates (Figure S1). However, where macrophages were present alongside bacteriophages, densities were detectable and thereby significantly higher (𝑥̅ = 1.3, 95%CI = 0.921 - 1.683) than where only bacteriophages were present (ANOVA comparing models with and without macrophage x bacteriophage presence interaction: 𝑥^"^ = 14.32, p = 0.001; Tukey HSD: t-ratio = -5.04, p < 0.001; Figure S1). Comparisons between the bacteria-only control and bacteriophage alone (Tukey HSD: t-ratio = 21.01, p < 0.001) and macrophage-bacteriophage treatment (Tukey HSD: t-ratio = 15.97, p < 0.001) were significant (Figure S1). Macrophages alone did not significantly decrease bacterial densities compared to controls (Tukey HSD: t-ratio = 0.31, p = 0.989; Figure S1).

This result indicated antagonism between bacteriophages and macrophages in how they affect bacteria density. We considered whether macrophages directly degraded bacteriophages by culturing bacteriophages alone with macrophages (no bacteria present). Macrophages were not found to significantly affect bacteriophage densities over 8 hrs of culturing (F_1,9_ = 0.044, p = 0.839; Figure S2).

### Bacteria and bacteriophage densities during experimental evolution

We next considered how macrophages shape bacteria-phage dynamics over timescales where (co)evolution may play an important role (six days; a treatment relevant timeframe) (Figure S3). Here, *P. aeruginosa* was transferred daily to fresh media with fresh macrophages in the macrophage-present treatments. Bacteriophages (14-1 and PNM) were inoculated at the start of the experiment and transferred with bacteria, with additional daily inoculations of fresh bacteriophage to mimic repeated bacteriophage dosing as done in phage therapy (Figure S3).

On their own, macrophages (ANOVA comparing models with and without macrophage x time interaction: 𝑥^"^ = 42.1, p < 0.001; Table S1) and bacteriophages (ANOVA comparing models with and without bacteriophage x time interaction: 𝑥^"^ = 25.86, p < 0.001; Table S2) decreased bacterial densities relative to controls (Figure 1a). Bacteriophages resulted in the greatest decrease (41.62%) in bacterial density compared to macrophages (8.73%) when averaged through time.

**Figure 1.**
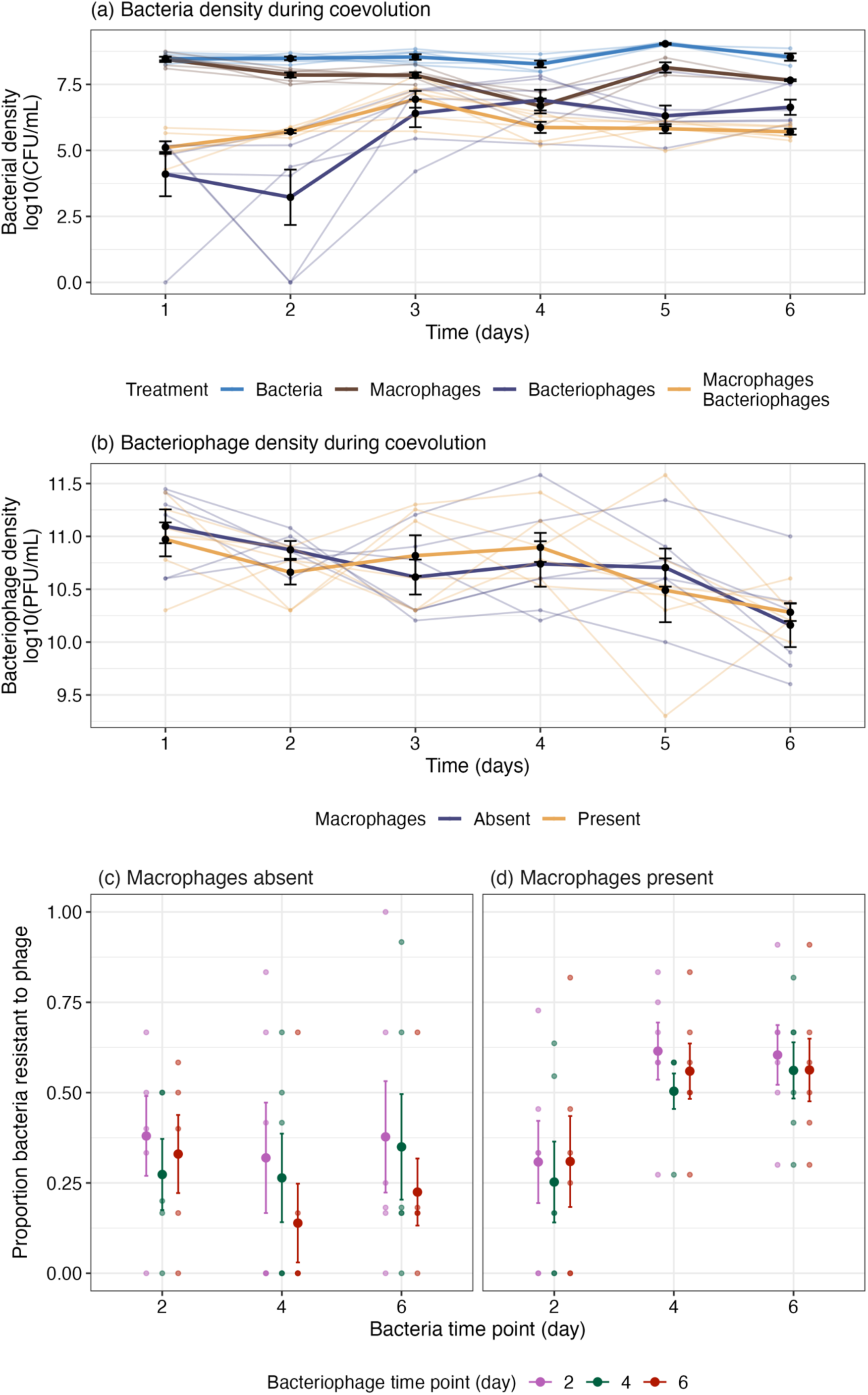
Ecological and evolutionary dynamics observed over experimental evolution. (a) Bacterial density (log10(CFU/mL)) in the presence and absence of bacteriophages and macrophages; and (b) bacteriophage density (log10(PFU/mL)) in the presence and absence of macrophages. Lines connect individual treatment replicates and changes in the treatment mean over time. Subsequently, a time-series assay was performed to analyse the proportion of bacteria resistant to a population of bacteriophages from different time-points where macrophages are (c) present and (d) absent. Small points indicate individual treatment replicates. Large points show the treatment mean. Bars indicate standard error (± SE).

We next considered results from the entire experiment analysed together. Here, we found a three-way interaction between the effects of macrophage and bacteriophage presence and the day of the experiment (ANOVA comparing models with and without the three-way interaction between time, macrophage and bacteriophage presence: 𝑥^"^ = 13.34, p = 0.0204). On day one, bacteriophages primarily decreased bacterial densities, with densities lower than controls and macrophage-only treatments (Tukey HSD < 0.05; Table S3; Figure 1a). Macrophages alone had no significant affect on bacteria densities relative to controls (Tukey HSD: p > 0.05; Table S3), except at day 4 (Tukey HSD: estimate = 1.59, t-ratio = 2.67, p-value = 0.043). Crucially, macrophages and bacteriophages interacted antagonistically (i.e. less than their additive effects) with respect to reducing bacterial densities. There was a notable increase in bacterial density with macrophages and bacteriophages present at day two (𝑥̅ = 5.71, 95%CI = 4.99 - 6.43), compared to the bacteriophage-only treatment (𝑥̅ = 3.23, 95%CI = 2.51 - 3.94; Tukey HSD: t-ratio = -4.84, p < 0.001). Day three onwards, bacterial densities with bacteriophage increased above that of the macrophage-bacteriophage treatment, although comparisons were non-significant (Tukey HSD: > 0.05; Table S3). Overall, the key results over 6 days show a greater effect of bacteriophage than macrophages, and antagonism between them which qualitatively matches the findings of antagonism between bacteriophages and macrophages over the 8 hour experiment.

Bacteriophage density was not significantly affected by macrophages (ANOVA comparing models with and without macrophage x time: 𝑥^"^ = 3.68, p = 0.596; fixed effect of macrophage presence: 𝑥^"^ = 0.01, p = 0.939) and were stable across most time-points (Tukey HSD comparisons between time-points 1 - 5: p > 0.05; Table S4; Figure 1b). However, by day six, bacteriophage densities showed significant decline in both treatments (Tukey HSD comparisons between time-points 1-5 and 6: p < 0.05; Table S4; Figure 1b).

### Bacteria-phage coevolution

We first determined the extent of bacterial resistance evolution to the two ancestral bacteriophages, 14-1 and PNM at the end of the experiment (day 6), with the ancestral bacteria having no resistance to these bacteriophages. Interestingly, no bacteriophage resistance emerged to PNM in all macrophage replicates (Figure S4). In the absence of macrophages, PNM resistance occurred in 2/6 treatment replicates at day six (16.7% and 8.3%). Bacteriophage resistance to 14-1 was non-significantly different between macrophage-present cultures (56.3%, 7.6 ± SE) compared to when bacteria were cultured with bacteriophages alone (30.6%, 14.5 ± SE; F_1,10_ = 1.51, p = 0.247; Figure S4).

We next considered if bacteria and bacteriophage (co)evolution occurred using a time-shift assay in which bacteria from different time-points are exposed to bacteriophages from different time-points (Figure S3). In the absence of macrophages, 32.8% (5.8 ± SE) of bacteria isolates were bacteriophage resistant by day two and this did not significantly increase through time (day 4 = 24.1%, 7.3 ± SE; day 6 = 31.7%, 7.4 ± SE; ANOVA comparing models with and without bacteriophage x bacteria time interaction: 𝑥^"^ = 4.44, p = 0.350; ANOVA comparing models with and without bacterial time-point: 𝑥^"^ = 4.64, p = 0.098; Figure 1b). Comparatively, the bacteriophage cocktail significantly increased in infectivity through time (ANOVA comparing models with and without bacteriophage time: 𝑥^"^ = 9.76, p = 0.008; Tukey HSD of bacteriophage infectivity between days two and six: z-ratio = 3.073, p = 0.006). This pattern is broadly indicative of asymmetrical coevolution with increases in bacteriophage infectivity occurring without increases in bacterial resistance.

In contrast, where macrophages were present, the proportion of bacteriophage resistant bacteria increased through time from 29% (6.4 ± SE) resistant at day 2 to 57.6% (8.2 ± SE) resistant at day six (ANOVA comparing models with and without bacterial time-point: 𝑥^"^ = 48.51, p < 0.001; Tukey HSD: estimate = -1.301, z-ratio = - 6.05, p < 0.001). Unlike the no-macrophage treatment, the bacteriophage cocktail did not increase in infectivity through time (ANOVA comparing models with and without bacteriophage x bacteria time interaction: 𝑥^"^ = 0.662, p = 0.956; ANOVA comparing models with and without bacteriophage time: 𝑥^"^ = 2.45, p = 0.294; Figure 1c). Therefore, the presence of macrophages limited coevolution with only increases in the overall rates of bacteriophage resistance without changes to bacteriophage infectivity.

These results demonstrate macrophages affected patterns of bacteriophage resistance and changes in bacteriophage cocktail efficacy. These different patterns led to differences in contemporary resistance i.e. the resistance of bacteria to the population of bacteriophages isolated from the same time point (e.g. resistance of bacteria from day six to bacteriophages from day six). Where macrophages were absent, contemporary bacteriophage resistance was significantly lower (day four: 𝑥̅ = 0.27, 95%CI = 0.164 - 0.420; day six: 𝑥̅ = 0.22, 95%CI = 0.125 -0.353) compared to where macrophages were present at days four (macrophages absent: 𝑥̅ = 0.27, 95%CI = 0.164 - 0.420; macrophages present: 𝑥̅ = 0.51, 95%CI = 0.363 - 0.650, Tukey HSD: estimate = 1.004, z-ratio = 2.24, p = 0.025) and six (macrophages absent: 𝑥̅ = 0.22, 95%CI = 0.125 - 0.353; macrophages present: 𝑥̅ = 0.57, 95%CI = 0.422 - 0.708, Tukey HSD: estimate = 1.56, z-ratio = 3.40, p < 0.001; ANOVA comparing models with and without treatment x time interaction: 𝑥^"^ = 13.48, p = 0.001; Figure 1d). This effect was not present at day two, however this was before bacteria in macrophage-present cultures increased in bacteriophage resistance, and before the bacteriophage cocktail increased in infectivity in the macrophage-absent cultures (Tukey HSD: estimate = - 0.398, z-ratio = -0.85, p = 0.395). Consequently, bacteria were overall more resistant to bacteriophages when macrophages were present and macrophages stopped increases in bacteriophage cocktail efficacy through time.

### Genetic changes within bacteria and bacteriophage populations

Sequencing analyses were conducted to determine the genetic underpinnings of bacteria-phage (co)evolution (Figure S3). Interestingly, few mutations were found in bacteria populations (Figure S5). No mutations were found in bacteria-only and bacteria-macrophage (no bacteriophage) treatments. In treatments containing bacteriophages, mutations were found in three out of six replicates of each treatment (six total, macrophage present and absent). In four out of six of these populations, 1 missense or frameshift mutation was found in a hypothetical protein upstream of a gene associated with LPS biosynthesis (Figure S5; Supplementary File 1). 14-1 infects via LPS therefore it is likely this mutation is associated with 14-1 resistance. Only one population had more than one mutation (three) including in genes associated with pili maturation and two hypothetical proteins (one in a region of chloramphenicol resistance genes and the other upstream of LPS biosynthesis) – this population was the only treatment replicate in which resistance to 14-1 reached fixation suggesting multiple mutations were required for this to occur (Figure S6). The remaining bacterial populations had mutations at a hypothetical protein associated with a chloramphenicol resistance region (upstream and downstream genes) and dihydrootase which are responsible for pyrimidines nucleotide synthesis – the mechanistic link between these mutations and bacteriophage resistance is unclear (Figure S6). Additionally, some populations which had bacteriophage resistant isolates had no genetic mutations. This suggests that resistance also occurred via mechanisms such as phase variation in which the expression of genes, such as for bacteriophage receptors, is turned off (31, 32). These alterations are heritable but are not identifiable via nucleotide sequencing (31, 32).

Next, we considered the bacteriophage populations. Overall, our sequencing analysis of PNM populations revealed no mutations compared to the ancestor while 15 and 24 mutations were found 14-1 populations evolved in the absence and presence of macrophages respectively. Most mutations (36/39) were missense mutations in bacteriophage tail fibre proteins while the remaining mutations (3/39) were in DNA helicase (missense mutation) and two hypothetical proteins (one missense, one conservative in-frame deletion; Supplementary File 2). Time-shift assays of coevolution suggested that bacteriophages increased in infectivity where macrophages were absent. If this effect was driven by genetic adaptation in the bacteriophage, we would expect more mutations in the macrophage-absent populations. Contrary to our predictions, 14-1 evolved significantly more in the presence of macrophages (𝑥̅ = 334, 95% = 248.8 – 418) compared to when macrophages were absent (𝑥̅ = 178, 95% CI = 93.5 – 263; ANOVA comparing models with and without macrophage presence: 𝑥^"^ = 8.35, p = 0.016; Tukey HSD comparing genetic distance between bacteriophage populations from macrophage absent to present cultures: estimate = -155, t-ratio = -2.89, p = 0.016; Figure 2).

**Figure 2.**
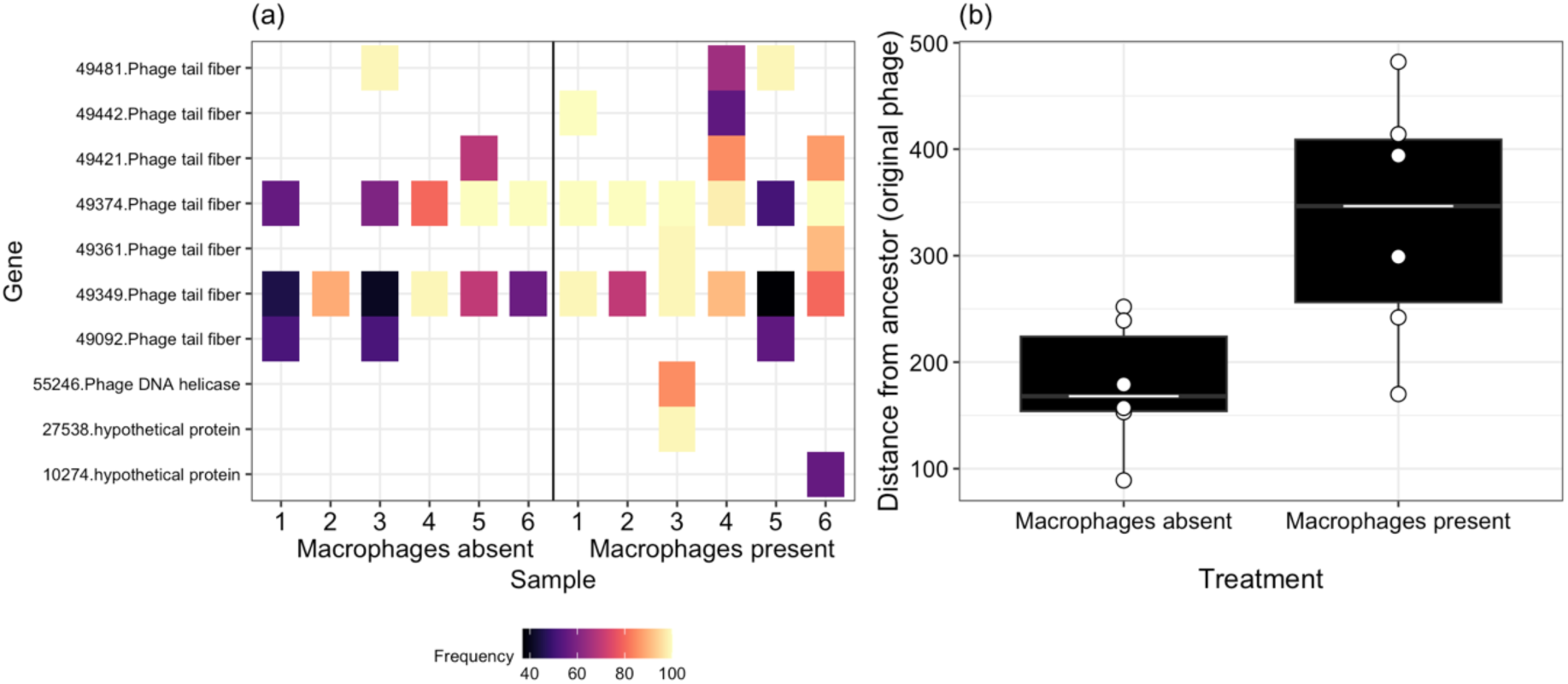
Bacteriophage (14–1) evolution in the presence and absent of macrophages at day 6. (a) In the presence of macrophages, a greater number of unique genetic variants are selected and increase to higher frequencies (%) within the population. On the y-axis the number indicates the position of the genetic variant within the genome while the gene name indicates the type of gene the mutation occurred in. 1-6 on the x-axis indicates the replicate number of each treatment. (b) These evolutionary differences can be summarised into the overall distance from the ancestor (sum of genetic variants and their frequencies) which shows that bacteriophages diverge more from the ancestor when macrophages are present. Points indicate individual treatment replicates. Tops and bottoms of the bars represent the 75th and 25th percentiles of the data, the middle lines are the medians, and the whiskers extend from their respective hinge to the smallest or largest value no further than 1.5* interquartile range.

Differences in bacteriophage resistance and patterns of coevolution may be in-part explained by differences in PNM frequences between macrophage present and absent cultures, in addition to any evolutionary change. Sequencing showed that PNM was present in only 1/6 macrophage-present replicates, while 4/6 treatment replicates contained PNM when macrophages were absent. The absence of PNM in macrophage-present cultures explains why there was no resistance evolution to this bacteriophage in these replicates, while resistance to PNM emerged where macrophages were absent in 2/6 treatment replicates. Additionally, where macrophages were absent, the percentage of reads mapping to PNM increased from 28.3% (6.65 ± SE) at day two to 35.01% (12.2 ± SE) at day six. Although the increase was statistically non-significant (ANOVA comparing models with and without time: 𝑥^"^ = 0.718, p = 0.397), 4/6 treatment replicates showed an increase in PNM proportion between days 2 and 6. Therefore, considering that most bacteria remained susceptible to PNM, it is likely that changes in PNM frequency also contributed to increases in bacteriophage cocktail infectivity. Since there was no statistical relationship between the frequency of bacteriophage resistant bacteria and proportion of PNM at each time point (ANOVA comparing models with and without PNM proportion: 𝑥^"^ = 1.01, p = 0.316), there are likely to be other variables influencing bacteriophage cocktail infectivity through time.

### Bacteriophages growth rates in the presence of macrophages

We next explored why 14-1 might have outcompeted PNM in the presence but not absence of macrophages. Macrophages may have reduced the relative growth rate of PNM by directly inhibiting contact rates between hosts and bacteriophage, or indirectly by altering the environmental media with the release of metabolites. We competed 14-1 and PNM in normal media (Dulbecco’s Modified Eagle’s Medium), spent macrophage media (from non-activated macrophages), media from activated macrophages (stimulated with heat-killed *P. aeruginosa*), and in fresh media with macrophages present (as set up in the original evolution experiment). Neither macrophages nor media type significantly affected the relative fitness between the bacteriophages when in competition (F_3,16_ = 1.95, p = 0.162, Figure S6). However, the presence of macrophages did significantly decrease the growth rates of the bacteriophages (ANOVA comparing models with and without treatment: 𝑥^"^= 57.97, p < 0.001; Tukey HSD comparisons: p < 0.001, Table S5, Figure 2) suggesting that macrophages can inhibit bacteria-phage contact rates to some extent. This effect is greater, but consistent with the non-significant trend in bacteriophage densities observed in the first two days of the coevolution experiment. Surprisingly, the growth rate of 14-1 (𝑥̅ = 14.3 *m*, 95% CI = 14.2 – 14.5) was on average lower than PNM (𝑥̅ = 14.6 *m*, 95% CI = 14.4 – 14.7; ANOVA comparing models with and without bacteriophage identity: 𝑥^"^ = 7.71, p = 0.005; Tukey HSD comparing growth rates of 14-1 to PNM: estimate = -0.25, t-ratio = -2.73, p = 0.013), however this difference was not significantly affected by treatment (ANOVA comparing models with and without bacteriophage identity and treatment interaction: 𝑥^"^ = 7.3, p = 0.063; Figure 3).

**Figure 3.**
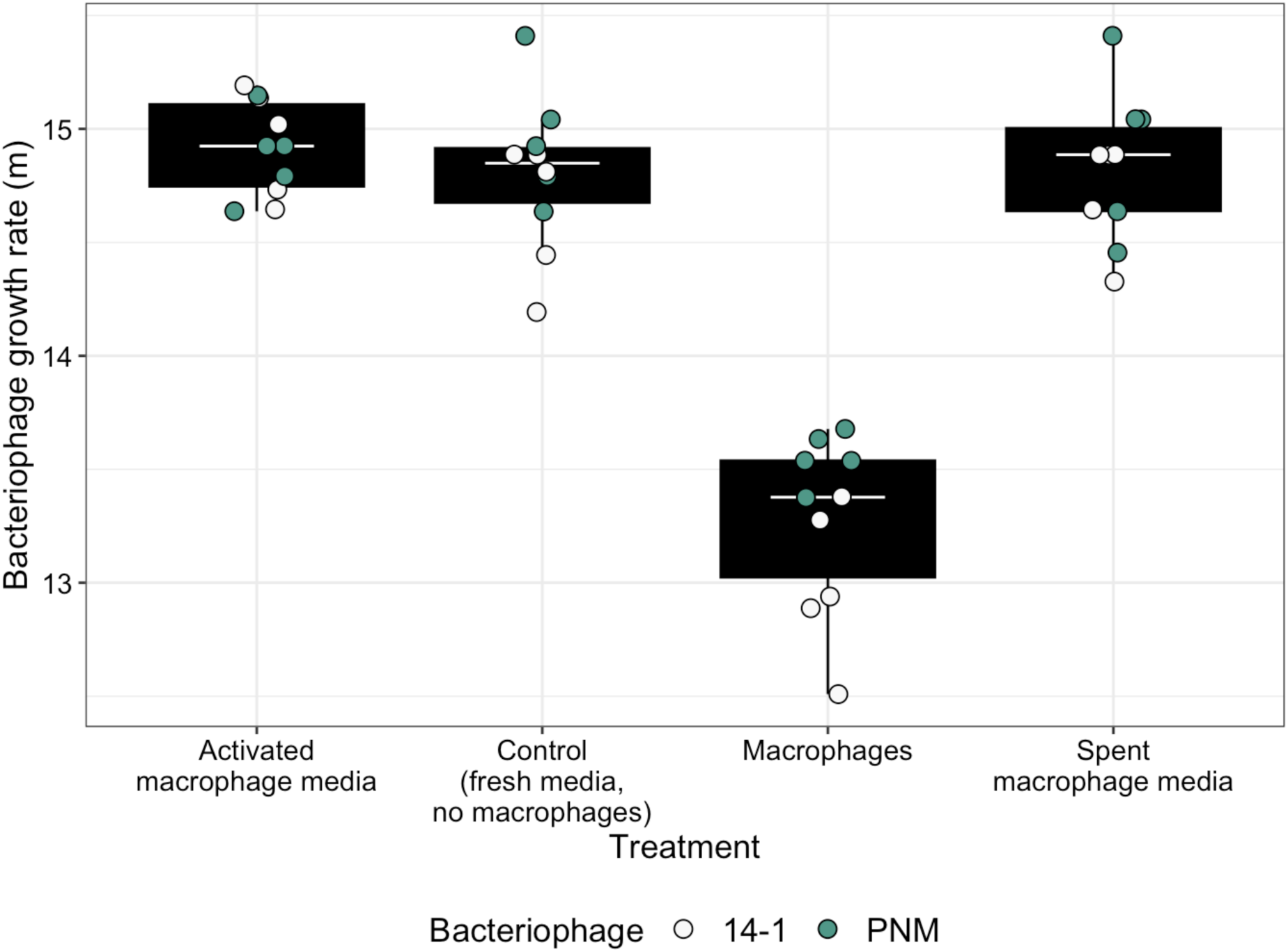
Bacteriophage (phage) growth rates (m) in different media and macrophage treatments. Tops and bottoms of the bars represent the 75th and 25th percentiles of the data, the middle lines are the medians, and the whiskers extend from their respective hinge to the smallest or largest value no further than 1.5* interquartile range.

### Effect of bacteriophage resistance on biofilm production and inflammatory cytokine production

Finally, we considered the consequences of bacteriophage resistance on biofilm formation and macrophage stimulation (Figure S3). On average, bacteriophage resistant populations produced significantly less biofilm (𝑥̅ = 6.22 log(fluorescence), 95%CI = 6.08 - 6.37) than bacteriophage susceptible populations (𝑥̅ = 6.43, 95%CI = 6.36 - 6.51; ANOVA comparing models without and without bacteriophage resistance: F_1, 15_ = 7.58, p = 0.015; Tukey HSD: estimate = 0.21, t-ratio = 2.75, p = 0.015; Figure 4a). Furthermore, bacteria evolved with macrophages and bacteriophages produced less biofilm (𝑥̅ = 6.21, 95%CI = 6.09 - 6.32) than bacteria evolved with bacteriophages but no macrophages (𝑥̅ = 6.5, 95%CI = 6.38 - 6.61; ANOVA comparing models with and without treatment: F_4, 15_ = 4.07, p = 0.02; Tukey HSD: estimate = -0.289, t-ratio = -3.77, p = 0.014; Figure 4a); suggesting some population phenotypic divergence under bacteriophage selection. All other comparisons between treatment groups were non-significant (Table S6). Differences in biofilm production were not due to growth rate costs associated with resistance (ANOVA comparing models with and without resistance x treatment interaction: F_1, 14_ = 1.15, p = 0.302; and resistance: F_1, 19_ = 0.819, p = 0.377) or evolution treatments (ANOVA comparing models with and without treatment: F_4, 15_ = 0.454, p = 0.769; Figure 4b); with no correlation between growth rate and biofilm production (F_1, 19_ = 2.23, p = 0.152).

**Figure 4.**
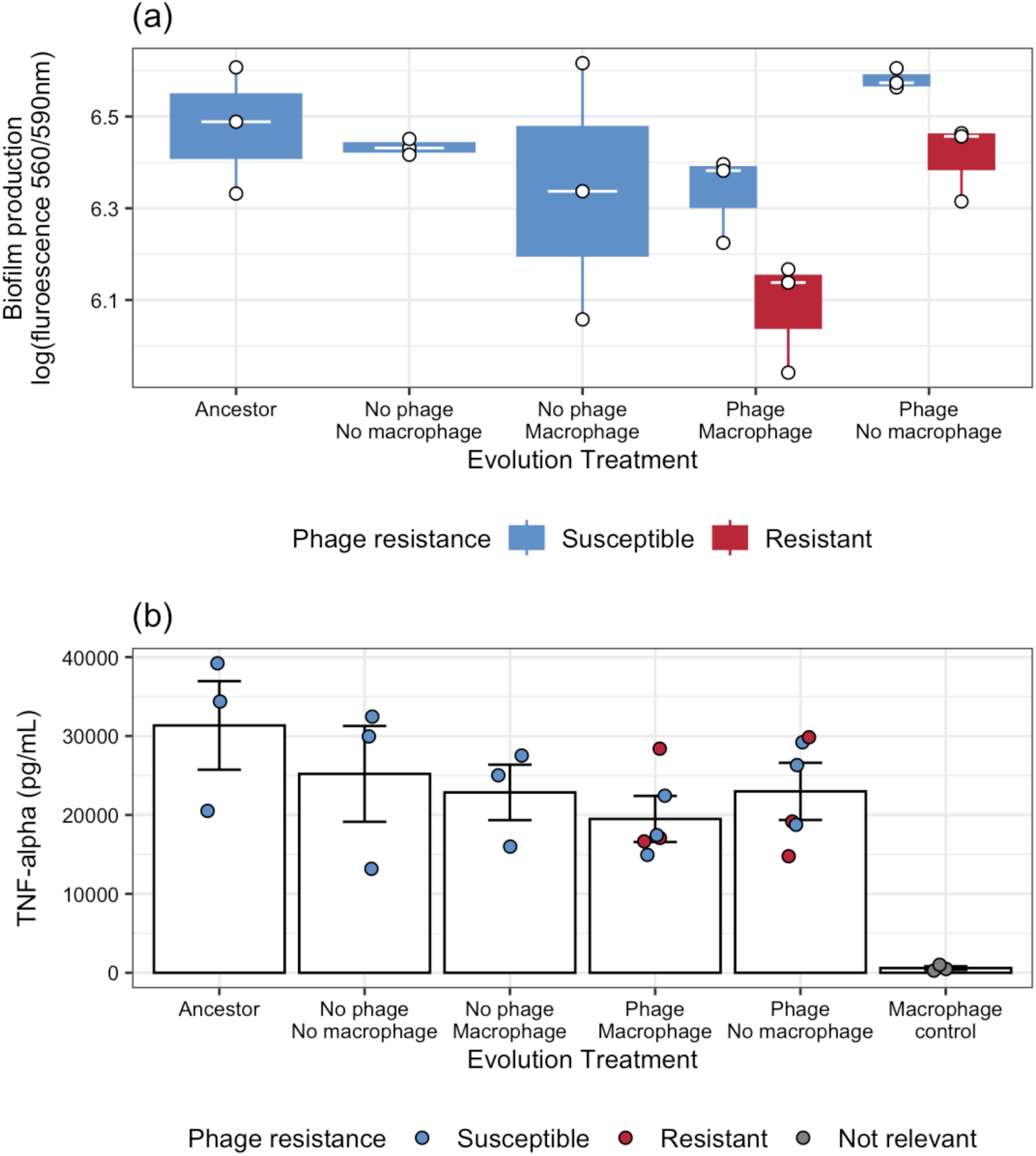
Phenotypic changes in bacteria associated with bacteriophage (phage) resistance and treatment bacteria evolved in during experimental evolution including (a) biofilm production and (b) macrophage stimulation, measured in production of pro-inflammatory cytokine production (TNF-α); bars indicate the treatment mean while error bars represent ± SE. Points indicate individual treatment replicates. Tops and bottoms of the bars represent the 75th and 25th percentiles of the data, the middle lines are the medians, and the whiskers extend from their respective hinge to the smallest or largest value no further than 1.5* interquartile range.

Levels of pro-inflammatory cytokine production (TNF-α) did not differ between bacteriophage resistant or susceptible isolates (ANOVA comparing models with and without bacteriophage resistance: F_1, 9_ = 0.05, p = 0.829) or between isolates evolved with bacteriophages and with or without macrophages (ANOVA comparing models without and without evolution with macrophages: F_1,10_ = 1.06, p = 0.327; and the interaction: F_1,8_ = 0.794, p = 0.399; Figure 4b). While there was a significant effect of treatment on cytokine production (ANOVA comparing models with and without evolution treatment: F_5,18_ = 57.2, p < 0.001), significant comparisons were only found between bacteria-present cultures and the macrophage-only control (no bacteria present) (Table S7). We also wanted to consider rates of phagocytosis, however due to the bacteria’s mucoid state it quickly forms biofilms resistant to antibiotic mediated killing, making it difficult to eliminate extracellular bacteria. However, based on these results and the strains’ resilience to macrophage killing (Figure S1), we do not expect this to differ by bacteriophage resistance or treatment.

## Discussion

How the immune system interacts with bacteriophage during treatment is hypothesised to have significant implications for treatment outcomes, but evidence for whether interactions are synergistic or antagonistic are mixed (11, 17, 18, 33). Here, our results suggest that interactions between bacteriophages and macrophages can be antagonistic over treatment relevant timescales. In the short term, macrophages limited bacteriophage growth rates which led to increased bacterial densities. Over longer timescales, macrophages increased the rate of bacteriophage resistance. Consequently, macrophages may have ecological and evolutionary consequences phage therapy efficacy.

Over short timescales (8 hrs and 1-2 days of experimental evolution), macrophages reduced the rate by which bacteria were killed by bacteriophages. Our growth rate assays with bacteriophages show that bacteriophage growth rates are inhibited when macrophages are present. This is consistent with recent work conducted in mice (25), although the mechanism appears different in our study. Since this effect is dependent on the macrophages being physically present (vs. media changes) while not directly degrading bacteriophages (as in (25)), we hypothesise that macrophages inhibit bacteria-phage contact rates. While typically considered an extracellular pathogen, *P. aeruginosa* has been shown to have an intracellular lifestyle and can survive after phagocytosis (34). As we observed bacteria inside our macrophages (observational results), but negligible effects of macrophages on bacteria density, we hypothesise that our *P. aeruginosa* strain CN573 is also resistant to killing once inside phagocytes.

Over longer timescales, we tracked changes in bacteriophage resistance, finding that macrophages increased overall rates of resistance. No resistance was observed to one of the bacteriophages in the cocktail, PNM, in most cultures, with resistance emerging in only 2/6 macrophage absent cultures, and at low frequency (<17%). While rates of resistance to the other ancestral bacteriophage in the cocktail, 14-1, were non-significantly higher when macrophages were present, a more clinically relevant measure of bacteriophage resistance is contemporary resistance – the levels of bacteriophage resistance of bacteria to bacteriophage from its own time-point Here, macrophages significantly increased bacteriophage resistance to contemporary bacteriophage from days four to six of experimental evolution. By decreasing the growth rates of bacteriophage, macrophages gave the bacteria an advantage early in the experiment likely which significantly increased bacterial mutation (including epigenetic potential) supply rates (13). As bacterial densities were very low, at almost undetectable densities, when just bacteriophages were present with bacteria, mutation supply rates for bacteriophage resistance were lower. This is synonymous to antibiotic resistance where incomplete treatment regimes allow bacteria to persist and evolve resistance. Furthermore, macrophages excluded one bacteriophage (PNM) from the cocktail, whereas both bacteriophages were present in 4/6 treatment replicates where macrophages were absent. As resistance to PNM was absent or relatively low, bacteria would have been significantly more susceptible to the contemporary bacteriophage where it was present (in the absence of macrophages). Additionally, presence of both bacteriophages may have constrained adaptation of the bacteria by resulting in reduced bacteria populations (in the short term). Adapting to two bacteriophage, versus one, is typically more costly to bacterial growth, which can further constrain expansion of bacteriophage resistant populations (35, 36). In our study, we were not able to define one effect of the macrophages on the bacteria or bacteriophage which resulted in increased bacteriophage resistant populations; however, this was likely a combination of greater bacterial densities early in the experiment due to macrophages constraining bacteriophage growth, and differences in bacteriophage cocktail diversity between treatments.

Mechanisms of bacteriophage resistance were unclear but appeared to be non-significantly different between cultures where macrophages were present and absent with overall similar resistance mutations. 4/12 treatment replicates (2 each for macrophage present/absent treatments) had mutations upstream of LPS which is likely associated with resistance to 14-1 (LPS binding). One macrophage-absent treatment replicate also had mutations in an intergenic region upstream of PilC – while PNM is pilus-binding, no resistance to PNM was found in this replicate, suggesting this may be a more general resistance mutation to 14-1. Few mutations were found, with many replicates that had bacteriophage resistant isolates having no detectable mutations – this raises the possibility that bacteriophage resistance also emerged by phase variation (32) which was undetectable in our sequencing analysis.

When examining patterns of coevolution, we observed an increase in bacteriophage infectivity when macrophages were absent between days 2 to 6, which was not present when macrophages were present. As with resistance patterns, we were not able to find one cause of increasing infectivity of the cocktail. However, a major driver was likely increases in PNM frequency which were only observed in the absence of macrophages. As resistance to PNM was relatively rare (although present in 2/6 treatment replicates), increases in PNM proportions would have resulted in more bacteria being susceptible to the bacteriophage at day six. Although the relationship between proportion of PNM in culture and susceptibility to bacteriophage from days two and six was non-significant, this may also be confounded by the emergence of resistance in the 2/6 replicates which were also the replicates where PNM was most frequent. Patterns of coevolution cannot be explained by genetic changes in the bacteriophages. No mutations were found in PNM and significantly more 14-1 mutations were found in bacteriophage where macrophages were present (and patterns of coevolution were absent). 14-1 mutation rates may have been driven by the higher rates of bacteriophage resistance in those cultures and/or adaptation to macrophages reducing bacteriophage growth rates. However, as bacteriophage densities were declining by day six, these mutations appear to have limited adaptive potential.

Although increases in bacteriophage resistance by macrophages is undesirable for bacteriophage efficacy, this can have beneficial effects with respect to reduced bacteria growth rates, biofilm production, antibiotic resistance and virulence (37). Here, bacteriophage resistance decreased biofilm production – this effect was independent of the presence of macrophages or any changes to growth rates. Decreases in biofilm production is a favourable outcome as bacteria typically become easier to clear by the immune system and antibiotics (23, 38). However, we did not find evidence that bacteriophage resistance increased susceptibility to macrophages, contrary to other studies (11, 17, 19). Furthermore, we did not see evidence of decreased proinflammatory cytokine production in macrophages exposed to bacteriophage resistant (lower biofilm producing) populations; this further emphasises that interactions with macrophages were not altered by bacteriophage resistance, which is likely to be very dependent on the mechanism of bacteriophage resistance (21). Although interactions with macrophages were not affected by bacteriophage resistance, the decreases in biofilm production is a beneficial outcome for infection management and patient prognosis in a phage therapy context (39).

Overall, these results suggest that macrophages can directly diminish bacteriophage cocktail efficacy by reducing bacteriophage growth rates and excluding one bacteriophage from the cocktail (reducing it to a monotherapy) which leads to higher rates of bacteriophage resistance. While bacteriophage cocktails are typically shown to be more effective than mono-therapies *in vitro* (40–43), their efficacy *in vivo* has been mixed with no strong evidence supporting their use over monotherapies (7, 44). The immune system, which will vary significantly between patients and infection type, likely has a key role mediating the success of bacteriophage cocktails. Consequently, future work should consider whether the bacteriophage efficacy, as typically measured without any host-associated variables, accurately reflects this efficacy *in vivo*. Comparisons of findings *in vivo*, in clinical cases, and *in vitro* may help to achieve this (22), alongside experiments with host-associated variables such as immune components and the microbiome.

This work adds to the growing body of work showing that the immune system can interfere with phage therapy. While we focus on macrophages, these cells are a key mediator of inflammation and the innate immune system (30). Future work will aim to place these findings into the wider context of the immune response including the innate and adaptive immune system. Different immune components, including neutrophils, complement and antibodies, may differentially affect bacteriophage efficacy against bacteria – therefore it will be important to determine the net effect of co-occurring immune components. This work importantly demonstrates the value of examining interactions between bacteria, bacteriophages and a component of the immune system over timescales beyond 24 hrs. Over longer timescales, bacteriophage cocktail composition can change, as demonstrated, and bacteriophages can evolve in response to bacteriophage resistance. How bacteriophages evolve during treatment is a unique concern to bacteriophage therapy, considering this treatment uses biological entities. Our work suggests the efficacy of bacteriophage cocktails should be tested in immune system contexts, with generalisations from laboratory studies treated with caution – this may aid the development of bacteriophage cocktails that work synergistically rather than antagonistically with the immune system to clear bacterial infections.

## Materials & Methods Cell line

Macrophages, RAW 264.7 cell lines (8 passages; lot number 70041657) were purchased from American Type Culture Collection (ATCC® TIB-71). Culture medium consisted of DMEM (Dulbecco’s Modified Eagle’s Medium, GlutaMAX) supplemented with 10% FBS (foetal bovine serum) and 1% penicillin/streptomycin (for subculturing only, not used in experiments with bacteria or bacteriophages). Cells were incubated at 5% CO_2_ and 90% humidity at 37°C. Cells were passaged at ∼70% confluency at a 1:10 seeding ratio in T-75 (75 cm^3^) flasks.

### Bacteria and bacteriophage strains

*Pseudomonas aeruginosa* strain CN573=PSE143 is a mucoid isolate which was selected based on its clinical origins (45, 46). Since *P. aeruginosa* strains can occupy clinical and natural environments, we selected a strain that would have a genetic background relevant for phage therapy research and susceptibility to clinically relevant bacteriophages. Bacteriophages included 14-1 and PNM, strains which are used together for phage therapy, most notably in the PhagoBurn clinical trial (47). This bacteriophage cocktail has previously been characterised to ensure no prophage elements are present or that the bacteriophage carry toxin-encoding genes that may increase bacterial resistance and/or virulence (46). Bacteriophages had been purified of endotoxins to prevent harm to eukaryotic cells (including the macrophages in this study) (46). As 14–1 infects via a lipopolysaccharide (LPS) (48, 49) receptor and PNM uses type IV pili (50), resistance to one bacteriophage is unlikely to result in cross-resistance and neither bacteriophage should compete for receptor sites (43). Bacteriophages were purified of endotoxins using Endotraps (Lionex) prior to experimental use. Endotoxins were measured using a Chromogenic Endotoxin Quantification Kit with target endotoxin concentrations <0.5 EU/mL.

### Macrophage-bacteriophage effects on bacterial density

We first aimed to validate how macrophages and bacteriophages interacted bacteria densities during the growth phase, and whether macrophages affected bacteriophage densities independently. *Pseudomonas aeruginosa* strain CN573=PSE143 was cultured overnight in 6 mL LB at 37°C. Macrophages were seeded into 6-well plates at a density of 7.5 x 10^5^ cells / well with 2.5 mL culture medium and incubated overnight to reach ∼80-90% confluency for experiments.

For bacteria experiments, four treatments were established: bacteria only, bacteria with macrophages, bacteria with bacteriophages, and bacteria with macrophages and bacteriophages. *P. aeruginosa* was diluted to ∼10^5^ CFU/ μL (0.1 ocular density (OD_600_; 600 nm wavelength)) into PBS (phosphate buffered saline) and 10 μL was inoculated into each microcosm (for macrophage cultures: ∼1.5:1 macrophage to bacteria). To cultures containing bacteriophage, 10^5^ PFUs of each bacteriophage were inoculated (1 μL of each bacteriophage stock at 10^5^ PFUs/μL) to achieve an MOI of 0.1 per bacteriophage (bacteria to bacteriophage ratio). For bacteriophage experiments, two treatments were established: bacteriophage only, and macrophages with bacteriophages. 10^7^ PFUs (50 μL of 2 x 10^5^ stock bacteriophage) was inoculated (for macrophage cultures: 1:10 macrophage to bacteriophage ratio) into each well. Plates were incubated in tissue culture incubators for 8 hrs. At the end of the experiment, cultures were mixed via pipetting and 500 μL was frozen with 500 μL 50% glycerol at -70 ℃. To estimate changes in bacterial density, cultures were plated from frozen onto LB agar and incubated overnight at 37°C. Bacteriophage extracted from the bacteriophage experiment via chloroform extraction: 900 μL of culture was vortexed with 100μL chloroform. Vials were then centrifuged at 14,000 rpm (21100 x g; Progen GenFuge 24D centrifuge) for five minutes and the supernatant isolated. Bacteriophage density was estimated via spot assays: dilutions were spotted onto overlays containing 100 μL of *P. aeruginosa* (grown overnight as above) in 5 mL soft LB agar.

### Experimental evolution

For experimental evolution, four treatments were established with six replicates per treatment: bacteria only, bacteria with macrophages, bacteria with bacteriophages, and bacteria with both macrophages and bacteriophages. Macrophages were seeded into T-25 (25 cm^3^) tissue culture flasks with 10^6^ cells and incubated overnight as above to reach ∼80-90% confluency. Cell culture medium was refreshed prior to inoculation. Overnights of *P. aeruginosa* (grown in Luria Broth (LB) at 37 °C, shaking at 180 r.p.m) were diluted to 0.1 OD and 20 μL was inoculated into each microcosm (2 x 10^6^ CFUs; ∼1:1 bacteria to macrophage ratio). To bacteriophage cultures, 40000 PFUs were inoculated (2 μL of bacteriophage stock at 20000 PFUs/uL) to achieve an MOI of 0.02 (bacteria to bacteriophage ratio). Flasks were incubated as above. Serial transfers took place every 24 hrs for six days. 24 hrs prior to transfer, fresh macrophages were seeded into flasks mimicking starting conditions with media refreshed. At each transfer, replicates of each evolution line were vortexed and 1 mL of culture was removed and centrifuged at 5000 rpm (2000 x g) (Progen GenFuge 24D centrifuge) for 5 minutes to pellet bacteria. The supernatant was removed and the pellet resuspended in PBS (phosphate-buffered saline) to reduce endotoxin transfer. 50 μL was transferred into fresh culture medium and at each transfer, 2 μL of bacteriophage stock was inoculated into relevant flasks. At each transfer, cultures were frozen (500 μL was frozen with 500 μL 50% glycerol at -70 °C) and bacteriophage extracted via chloroform extraction as above. Bacteria and bacteriophage densities were estimated as previous. No-bacteriophage controls were monitored for bacteriophage presence. 3 macrophage-only and 2 bacteria-only treatment replicates became contaminated during experimental evolution and were removed.

### Estimating coevolution

To estimate whether bacteria and bacteriophage have coevolved in isolation or in the presence of macrophages, we performed time-shift assays (16). From days two, four and six, twelve colonies were isolated from each treatment replicate. To remove bacteriophage contamination, single colonies were resuspended in 10 μL M9 and were serially streaked onto LB plates for three transfers. Colonies were then grown overnight at 37 ℃ and bacteriophage spot assays were performed to test resistance against bacteriophage isolates from days two, four and six. Bacteria isolates from day six were tested for resistance against ancestral bacteriophages 14-1 and PNM. If bacteria and bacteriophage have coevolved via arms-race dynamics, we would expect bacteria to be more resistant to bacteriophages from the past and more susceptible to bacteriophages from the future. Twelve colonies were isolated from each no-bacteriophage treatment replicate (bacteria alone and bacteria with macrophages day six) and tested for bacteriophage resistance to 14-1 and PNM.

### Sequencing

Ancestral bacteriophages 14-1 and PNM were sequenced to identify any mutations which may have occurred which distinguish our samples from the reference genome. Prior to DNA extraction, cultures were treated with DNase to remove bacteria DNA. The NEB monarch genomic DNA extraction kit was used to extract gDNA and the concentration was determined as above. Sequencing was done by the University of Liverpool’s Centre for Genomic Research (CGR) using the PacBio Sequel IIe platform to generate an estimated average of 10 Gb HiFi data per library. Genomes were assembled and annotated using BV-BRC (Bacterial and Viral Bioinformatics Resource Center) bioinformatic tools (v 3.46.3) using Canu (51). Genomes were annotated using the ‘bacteriophages’ tool kit with comparison to the reference genomes of 14-1 (taxon ID: 581037) and PNM (taxon ID: 2975222).

The *P. aeruginosa* CN573 strain was sent for sequencing to MicrobesNG who extracted gDNA, sequenced and assembled the genome using in-house protocols. Sequencing libraries were generated using SQK-RBK114.96. Sequencing was performed on a GridION (Oxford Nanopore Technologies) using an R10.4.1 flowcell, with basecalling model r1041_e82_400bps_sup_variant_v4.3.0. Reads were randomly subsampled to 50X coverage using Rasusa (V 0.7.1), and assembled using Flye (V2.9.2-b1786). Assemblies were polished using Medaka (V 1.11.3) and the relevant model (r1041_e82_400bps_sup_variant_v4.3.0). Genomes were annotated using BV-BRC annotation tools with comparison to the reference genome *P. aeruginosa* PAO1 (identified as the closest characterised *P. aeruginosa* genome).

Bacteria populations from each treatment replicate from day 6 were analysed for mutations. The 12 isolates isolated from each treatment replicate were pooled by mixing one colony of each isolate into extraction buffer. Pooled isolates were used to remove bacteriophage contamination which would have been present in original stocks taken from experimental evolution. Bacteriophage populations from days 2 and 6 were amplified from original stocks by inoculating 150 μL stock into 15 mL LB with 150 μL overnight culture of *P. aeruginosa* CN573. Cultures were grown overnight, shaking, at 180 r.p.m. Bacteriophage were extracted by filtering cultures through 0.22 μm syringe filters. Bacteriophages were amplified and treated with DNase as above. The Qiagen DNeasy Ultra Clean DNA Extraction Kit reagents and protocol were used to extract the gDNA from the cultures. The concentration of gDNA in each sample was determined using the QuBit dsDNA HS Assay kit regents and protocol. Sequencing was done by the University of Liverpool’s Centre for Genomic Research (CGR) using an Illumina Novaseq to create 150bp paired end reads. The CGR trimmed for the presence of Illumina adapter sequences using Cutadapt (v1.2.1) and removed reads with a minimum quality score of 20 and which were shorter 15bp in length. Variant analysis on day 6 populations was run using BV-BRC tools with bacteria and bacteriophages compared to the assembled genomes of the ancestral strains. BWA-mem was used to align reads and FreeBayes was used to identify high-quality variants. Prior to analysis, variants found in every read library were identified as sequencing errors and removed.

### Bacteriophage competition assay

We determined if macrophages affected the relative competition of the two bacteriophages in our cocktail: 14-1 and PNM. To determine whether effects were driven by macrophage presence (direct effect) or by macrophages altering the media (indirect effect), we generated spent media from macrophage sub-culturing and activated macrophage media. Activated macrophage media was generated by exposing macrophages (2 x 30 mL 175 cm^3^ flasks of macrophages at 90% confluency) to heat-killed *P. aeruginosa* CN573. *P. aeruginosa* was first grown overnight in LB broth before being pelleted and washes 2x in PBS as before remove metabolites and media. The pellet was resuspended in PBS and then vials were placed at 80°C for 30 minutes (time selected based on preliminary experiments). 1.8 mL of heat-inactivated bacteria was then added to each flask. Macrophages were incubated for 24 hrs as previous. After 24 hrs, media was removed and first spotted onto LB agar to confirm no bacteria contamination, before being filter sterilised using 0.45 μm and 0.22 μm syringe filters to remove cellular debris. Media was stored at -20 °C and thawed immediately prior to experimentation. For the macrophage-present treatment, macrophages were seeded into 6 25 cm^3^ flasks and grown overnight as previous to reach 90% confluency – media was refreshed prior to experimentation.

*P. aeruginosa* CN573 was cultured overnight, washed in PBS and diluted to 0.1 OD_600_ as previous. A master-mix of both bacteriophages was generated containing 1000 PFUs / μL of each bacteriophage. 20 μL of *P. aeruginosa* and 40 μL of bacteriophage master-mix was inoculated into independent flasks that contained either fresh DMEM, spent macrophage media, activated macrophage media, or macrophages. 5 independent replicates were set up for each treatment. Competition assays were run for 24 hrs. Samples were frozen and bacteriophage extracted via chloroform extraction as previous.

The density of each bacteriophage was determined by plating extractions onto lawns of three *P. aeruginosa* CN573 genotypes (wild-type (ancestor), PNM resistant, and 14-1 resistant). *P. aeruginosa* CN573 strains resistant to PNM and 14-1 were generated by inoculating 60 μL of *P. aeruginosa* into 6 mL LB with 10 μL stock of either 14-1 and PNM. Cultures were grown overnight before being streaked onto LB agar and grown overnight. Colonies were picked and resistance to each bacteriophage was confirmed by spot assays. Resistance to one bacteriophage did not affect plating efficiency of the other bacteriophage it was still susceptible to. The growth rate (m) of each bacteriophage was estimated as ln(N_1_/ N_0_) where N_1_ is the final density, N_0_ is starting density. The relative fitness of each bacteriophage was calculated by dividing the growth rate of 14-1 by the growth rate of PNM.

### Biofilm assay

We were interested whether resistance to bacteriophage affected bacteria phenotypes that could alter interactions with immune cells. As strain CN573 is mucoid, we considered whether bacteriophage resistance or evolution with macrophages had altered biofilm production – reductions in biofilm can make bacteria more vulnerable to the immune system. 6 bacteriophage resistant and 6 bacteriophage susceptible isolates was selected from independent treatment replicates from macrophage present and macrophage absent treatments (3 resistant/susceptible from each treatment). These were compared to 3 replicates of the ancestral bacteria, and 3 isolates from the bacteria-only evolution lines, and the bacteria-macrophage (no bacteriophage) evolution lines, all isolated from independent replicates. Biofilm production was estimated using a resazurin assay, based upon a previously described protocol (Peeters et al., 2008). Overnight monocultures were diluted to 0.1 OD_600_ (∼10^9^ CFU/ml) into PBS. 100 µl of each culture was inoculated into a sterile 96-well plate and incubated at 37℃ for 4 hr for cellular adhesion. Then, the supernatant was removed, and wells were washed gently with 100 µl of PBS by pipetting up and down three times. 100 µl of fresh LB media was added to each well including the controls (no bacteria) and plates were incubated for 16 hr. Liquid was removed and wells were washed with 100 µl PBS. 100 µl of fresh M9 was added followed by 20 µl Cell Titre Blue. Fluorescence (λex: 560 nm and λem: 590 nm) was measured after 2-hr incubation. Biofilm assays were independently replicated three times (technical replicates).

### Growth rate assay

To account for differences in bacteria growth rates in the biofilm assay, and to determine any costs of resistance, we measured exponential growth rates. The same bacteria isolates that were used in the biofilm assay were used. Overnight cultures were diluted to 0.1 OD_600_ into PBS. 10 µL of diluted cultured was inoculated into 190 µL LB media in 96-well plates. The plate was incubated in a spectrophotometer (MultiSkan Sky) at 37℃ in shaken conditions and OD_600_ was measured every 20 min for 24 hr to estimate changes in cellular density. Growth curves were independently replicated three times (technical replicates). Reads after 5 hrs of growth were removed as populations reached stationary phase with fluctuations in density estimates (likely due to biofilm formation) (Figure SX).

### Inflammatory cytokine production

We next assessed whether bacteriophage resistance led to changes in inflammatory cytokine production – specifically TNF-α as a key mediator of inflammation which is produced in response to bacteria infection. 6-well plates were seeded with 7.5 x 10^5^ macrophages and grown for 24 hrs to reach 90-100% confluency and media was refreshed. Bacteria were grown overnight (same isolates as used in the biofilm and growth rate assay) and diluted to 0.1 OD_600_ into PBS and 15 µL was inoculated into independent cultures (macrophage to bacteria ratio approx. 1:1). 3 macrophage cultures were not inoculated and kept as controls. Cultures were grown overnight in a tissue culture incubator as above. After culturing, 1 mL of culture was filtered through 0.45 μm and 0.22 μm syringe filters to remove bacteria and cellular debris.

Supernatants were stored at -80 °C and were thawed immediately prior to cytokine quantification. TNF-α concentrations were quantified using a ThermoFisher ELISA kit for mouse cytokines following manufacturer instructions. Samples were pre-diluted 100x so that values fit into the standard curve (identified in preliminary analysis). Samples were analysed in duplicate including blank controls – all results fit within 20% of the mean value. Absorbance OD_450_ and background absorbance OD_620_ was measured using spectrophotometer (MultiSkan Sky). A non-linear model was fitted to the standard curve of known TNF-α standard concentrations and resultant OD (after subtracting the background absorbance). TNF-α concentrations in the samples were estimated by predicting values based on the standard curve and multiplying by the dilution factor.

### Statistical analyses

All data were analysed using R (v.4.2.1) in RStudio (52), all plots were made using the package ‘*ggplot2*’ (53) and all tables were made using the package ‘*flextable*’ (54). Model simplification was conducted using likelihood ratio tests and Tukey’s post hoc multiple comparison tests were used to identify the most parsimonious model using the R package ‘*emmeans*’ (55). Bacterial density (after 8 hrs culturing) (log10(CFU/mL)) was analysed against interacting fixed effects of macrophage and bacteriophage presence in a linear model. Bacteriophage density after culturing with macrophages alone was determined in a linear model analysing bacteriophage density (log10(PFU/mL)) against macrophage presence. In experimental evolution, changes in bacterial density were analysed in linear mixed effects models. First, we were interested in the individual effects of macrophages and bacteriophages in reducing bacterial density relative to bacteria-only controls. As such, two models were construed analysing density (log10(CFU/mL)) against interacting fixed effects of time and macrophage or bacteriophage presence with a random effect of treatment replicate. In a final holistic model, density (log10(CFU/mL)) was tested against interacting fixed effects of macrophage presence, bacteriophage presence and time, with a random effect of treatment replicate. Changes in bacteriophage density was analysed in a linear mixed effects model analysing density (log10(PFU/mL)) against interacting fixed effects of time and macrophage presence with a random effect of treatment replicate.

Bacteriophage resistance to ancestral bacteriophage 14-1 on day 6 was analysed in a generalised linear model with a fixed effect of macrophage presence with a quasibinomial error structure. To assess whether bacteria and bacteriophage coevolved in the presence or absence of macrophages, a binomial generalised mixed effects linear model was used. Here, the proportion of bacteriophage resistant bacteria was analysed against interacting fixed effects of bacterial time and bacteriophage time with a random effect of treatment replicate. Separate models were constructed for each treatment (macrophages present or absent) to improve model fitting and primarily characterise whether coevolution occurred independent of treatment comparisons.

Bacteriophage populations were analysed based on the genetic distance from the ancestral population, calculated as the sum of the difference of the proportion of each SNP/indel in each population from the ancestral proportion. Genetic distance was analysed in a linear model with a fixed effect of macrophage presence. Changes in frequency of PNM in the bacteriophage cocktail was measured by analysing the proportion of reads that mapped to PNM (arcsine transformed) against a fixed effect of day (two or six) in a linear mixed effects model with a random effect of treatment replicate. The relationship between the proportion of bacteriophage resistant bacteria (taken as the average resistance to bacteriophage isolated from days two and six of the experiment) and proportion of PNM was analysed in a generalised linear model with a binomial error structure and a random effect of treatment replicate.

Competition assays between bacteriophages 14-1 and PNM were analysed in a linear model with relative fitness was analysed against treatment (media type / macrophage presence). Additionally, the bacteriophage growth rates (*m*) were analysed against interacting fixed effects of treatment and bacteriophage identity (14-1 or PNM) with a random effect of treatment replicate.

Biofilm production was log transformed (log(fluorescence λex: 560 nm and λem: 590 nm)) and analysed against interacting fixed effects of treatment (which evolution treatment clones originated from e.g. bacteria only, bacteria with macrophages etc.) and bacteriophage resistance. Growth rates were estimated using a rolling regression taking the steepest slope of the linear regression between ln OD_600_ and time in hours in a shifting window of every four time points (every 1.3 hr). Growth rate was averaged over each technical replicate for each biological replicate (individual bacterial isolates). Differences in growth rates were analysed against interacting fixed effects of treatment and bacteriophage resistance in a linear model. Additionally, biofilm production was analysed against growth rates in a linear model.

We analysed whether there were any significant differences in TNF-α cytokine production by macrophages when exposed to different bacteria populations. First, we analysed whether cytokine production (log transformed) differed across treatment groups including the macrophage control (unstimulated macrophages) in a linear model. Next, we assessed whether there were any differences in cytokine production between bacteriophage resistant and bacteriophage susceptible isolates originating from macrophage and bacteriophage treatment and bacteriophage only treatment. In a linear model, cytokine production was analysed against interacting fixed effects of macrophage and bacteriophage presence (during evolution).

## Author contributions

MC, SKW, JAW, GDB and AB conceived and designed the study. Experiments were conducted by MC, ZS, AM. DNA extractions were performed by MC, RL and SK. Data analysis conducted by MC. All authors contributed to the writing of the manuscript.

## Acknowledgements

We thank Stineke van Houte and Michael Brockhurst for helpful feedback on a manuscript draft; and Jean-Paul Pirnay and Maya Merabishvili for providing the endotoxin free bacteriophages. This work was funded by grant no. MR/N0137941/1 for the Great West 4 BIOMED Medical Research Council Doctoral Training Partnership, awarded to the Universities of Bath, Bristol, Cardiff, and Exeter from the Medical Research Council/UK Research and Innovation, awarded to MC. We acknowledge funding from the MRC Centre for Medical Mycology at the University of Exeter (MR/N006364/2 and MR/V033417/1), the NIHR Exeter Biomedical Research Centre. The views expressed are those of the author(s) and not necessarily those of the NIHR or the Department of Health and Social Care. This work was supported by NERC awards NE/V012347/1 and NE/S000771/1 awarded to AB.

## Data availability

R code and data will be deposited on GitHub.

