## Supplementary File 1 for "Antagonism between bacteriophages and macrophages decreases efficacy of a bacteriophage cocktail and increases bacteriophage resistance"

| Sample | Macrophages | Position | Score | Frequency | Type | Reference | Variant | Function | Upstream_feature | Downstream_feature | SNP type |
|---|---|---|---|---|---|---|---|---|---|---|---|
| BP1 | N | 504,704 | 100,699.0 | 0.90 | Nonsyn | agt | Cgt | hypothetical protein | hypothetical protein | probable lipopolysaccharide biosynthesis translocase NMA0643 | missense_variant |
| BMP2 | Y | 505,106 | 74,066.9 | 0.77 | Nonsyn | ttt | Gtt | hypothetical protein | hypothetical protein | probable lipopolysaccharide biosynthesis translocase NMA0643 | missense_variant |
| BP3 | N | 505,394 | 30,368.6 | 0.65 | Deletion | ggaggggggcat | GGAGGGGGCat | hypothetical protein | hypothetical protein | probable lipopolysaccharide biosynthesis translocase NMA0643 | frameshift_variant |
| BMP5 | Y | 505,394 | 41,616.8 | 0.65 | Deletion | ggaggggggcat | GGAGGGGGCat | hypothetical protein | hypothetical protein | probable lipopolysaccharide biosynthesis translocase NMA0643 | frameshift_variant |
| BP4 | N | 276,775 | 15,460.0 | 0.56 | Nonsyn | tac | Gac | Dihydroorotase (EC 3.5.2.3) | Aspartate carbamoyltransferase (EC 2.1.3.2) | Cystathionine gamma-lyase (EC 4.4.1.1) | missense_variant |
| BMP4 | Y | 843,757 | 78,036.7 | 0.76 | Nonsyn | gac | gaG | hypothetical protein | Chloramphenicol O-acetyltransferase (EC 2.3.1.28) | Chloramphenicol acetyltransferase (EC 2.3.1.28) | missense_variant |
| BP1 | N | 844,229 | 60,677.8 | 0.77 | Nonsyn | gtg | Ttg | hypothetical protein | Chloramphenicol O-acetyltransferase (EC 2.3.1.28) | Chloramphenicol acetyltransferase (EC 2.3.1.28) | missense_variant |
| BP1 | N | 114,112 | 119,715.0 | 1.00 | Nonsyn | tgg | tAg | Leader peptidase (Prepilin peptidase) (EC 3.4.23.43) / N-methyltransferase (EC 2.1.1.-) | Dephospho-CoA kinase (EC 2.7.1.24) | Type IV fimbrial assembly protein PilC | stop_gained |
