## Supplementary File 2 for "Antagonism between bacteriophages and macrophages decreases efficacy of a bacteriophage cocktail and increases bacteriophage resistance"

| Sample | Macrophages | Position | Frequency | Type | Reference | Variant | Function | SNP type |
|---|---|---|---|---|---|---|---|---|
| BMP1T6 | Y | 49349 | 0.99 | Nonsyn | aat | Tat | Phage tail fiber | missense_variant |
| BMP1T6 | Y | 49374 | 1.00 | Nonsyn | caa | cGa | Phage tail fiber | missense_variant |
| BMP1T6 | Y | 49442 | 1.00 | Nonsyn | caa | Aaa | Phage tail fiber | missense_variant |
| BMP2T6 | Y | 49349 | 0.70 | Nonsyn | aat | TAt | Phage tail fiber | missense_variant |
| BMP2T6 | Y | 49374 | 1.00 | Nonsyn | caa | cGa | Phage tail fiber | missense_variant |
| BMP3T6 | Y | 27538 | 0.99 | Nonsyn | gcc | gTc | hypothetical protein | missense_variant |
| BMP3T6 | Y | 49349 | 0.99 | Nonsyn | aat | CGt | Phage tail fiber | missense_variant |
| BMP3T6 | Y | 49361 | 0.99 | Nonsyn | atc | Ttc | Phage tail fiber | missense_variant |
| BMP3T6 | Y | 49374 | 1.00 | Nonsyn | caa | cGa | Phage tail fiber | missense_variant |
| BMP3T6 | Y | 55246 | 0.85 | Nonsyn | atg | Gtg | Phage DNA helicase | missense_variant |
| BMP4T6 | Y | 49349 | 0.91 | Nonsyn | aat | Tat | Phage tail fiber | missense_variant |
| BMP4T6 | Y | 49374 | 0.98 | Nonsyn | caa | cGa | Phage tail fiber | missense_variant |
| BMP4T6 | Y | 49421 | 0.85 | Nonsyn | gcc | Tcc | Phage tail fiber | missense_variant |
| BMP4T6 | Y | 49442 | 0.55 | Nonsyn | caa | Aaa | Phage tail fiber | missense_variant |
| BMP4T6 | Y | 49481 | 0.65 | Nonsyn | gtc | Ttc | Phage tail fiber | missense_variant |
| BMP5T6 | Y | 49092 | 0.55 | Nonsyn | gga | gTa | Phage tail fiber | missense_variant |
| BMP5T6 | Y | 49349 | 0.37 | Nonsyn | aat | CAt | Phage tail fiber | missense_variant |
| BMP5T6 | Y | 49374 | 0.51 | Nonsyn | caa | cGa | Phage tail fiber | missense_variant |
| BMP5T6 | Y | 49481 | 0.99 | Nonsyn | gtc | Ttc | Phage tail fiber | missense_variant |
| BMP6T6 | Y | 10274 | 0.56 | Deletion | tcgcagcggtcgcagcggtcc | tCGCAGCGGTCC | hypothetical protein | conservative_inframe_deletion |
| BMP6T6 | Y | 49349 | 0.80 | Nonsyn | aat | Tat | Phage tail fiber | missense_variant |
| BMP6T6 | Y | 49361 | 0.91 | Nonsyn | atc | Ttc | Phage tail fiber | missense_variant |
| BMP6T6 | Y | 49374 | 1.00 | Nonsyn | caa | cGa | Phage tail fiber | missense_variant |
| BMP6T6 | Y | 49421 | 0.87 | Nonsyn | gcc | Tcc | Phage tail fiber | missense_variant |
| BP1T6 | N | 49092 | 0.52 | Nonsyn | gga | gTa | Phage tail fiber | missense_variant |
| BP1T6 | N | 49349 | 0.45 | Nonsyn | aat | Tat | Phage tail fiber | missense_variant |
| BP1T6 | N | 49374 | 0.56 | Nonsyn | caa | cGa | Phage tail fiber | missense_variant |
| BP2T6 | N | 49349 | 0.89 | Nonsyn | aat | Tat | Phage tail fiber | missense_variant |

| Sample | Macrophages | Position | Frequency | Type | Reference | Variant | Function | SNP type |
|---|---|---|---|---|---|---|---|---|
| BP3T6 | N | 49092 | 0.52 | Nonsyn | gga | gTa | Phage tail fiber | missense_variant |
| BP3T6 | N | 49349 | 0.41 | Nonsyn | aat | CAt | Phage tail fiber | missense_variant |
| BP3T6 | N | 49374 | 0.60 | Nonsyn | caa | cGa | Phage tail fiber | missense_variant |
| BP3T6 | N | 49481 | 0.99 | Nonsyn | gtc | Ttc | Phage tail fiber | missense_variant |
| BP4T6 | N | 49349 | 0.99 | Nonsyn | aat | Tat | Phage tail fiber | missense_variant |
| BP4T6 | N | 49374 | 0.80 | Nonsyn | caa | cGa | Phage tail fiber | missense_variant |
| BP5T6 | N | 49349 | 0.70 | Nonsyn | aat | Tat | Phage tail fiber | missense_variant |
| BP5T6 | N | 49374 | 1.00 | Nonsyn | caa | cGa | Phage tail fiber | missense_variant |
| BP5T6 | N | 49421 | 0.69 | Nonsyn | gcc | Tcc | Phage tail fiber | missense_variant |
| BP6T6 | N | 49349 | 0.57 | Nonsyn | aat | CAt | Phage tail fiber | missense_variant |
| BP6T6 | N | 49374 | 1.00 | Nonsyn | caa | cGa | Phage tail fiber | missense_variant |
