## Supplementary material for "Antagonism between bacteriophages and macrophages decreases efficacy of a bacteriophage cocktail and increases bacteriophage resistance": Table S

Table S1. Multiple pairwise comparisons comparing bacterial densities when macrophages were absent (“N”) and present (“Y”) through days 1-6 of experimental evolution.

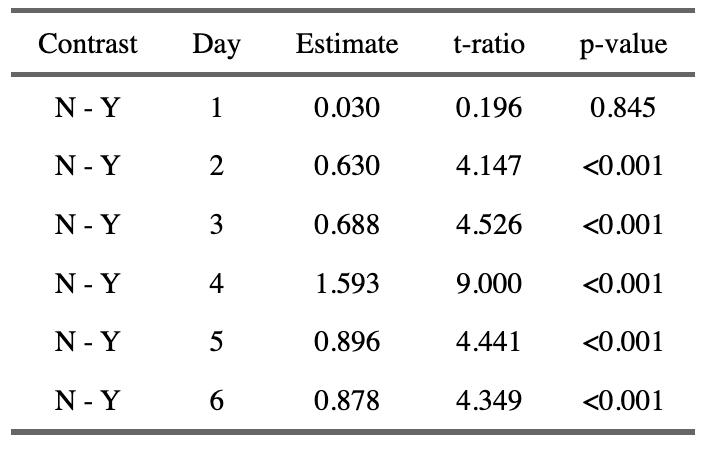

Table S2. Multiple pairwise comparisons comparing bacterial densities when bacteriophages were absent (“N”) and present (“Y”) through days 1-6 of experimental evolution.

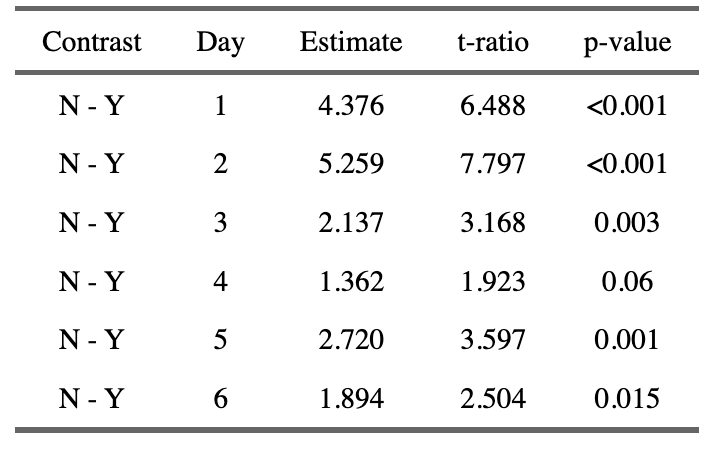

Table S3. Multiple pairwise comparisons comparing bacterial densities across four treatments where bacteriophages and macrophages were present (“Both” when together) and absent (“Bacteria”) through days one to six of experimental evolution. P-values adjusted using the Tukey method of comparing a family of four estimates.

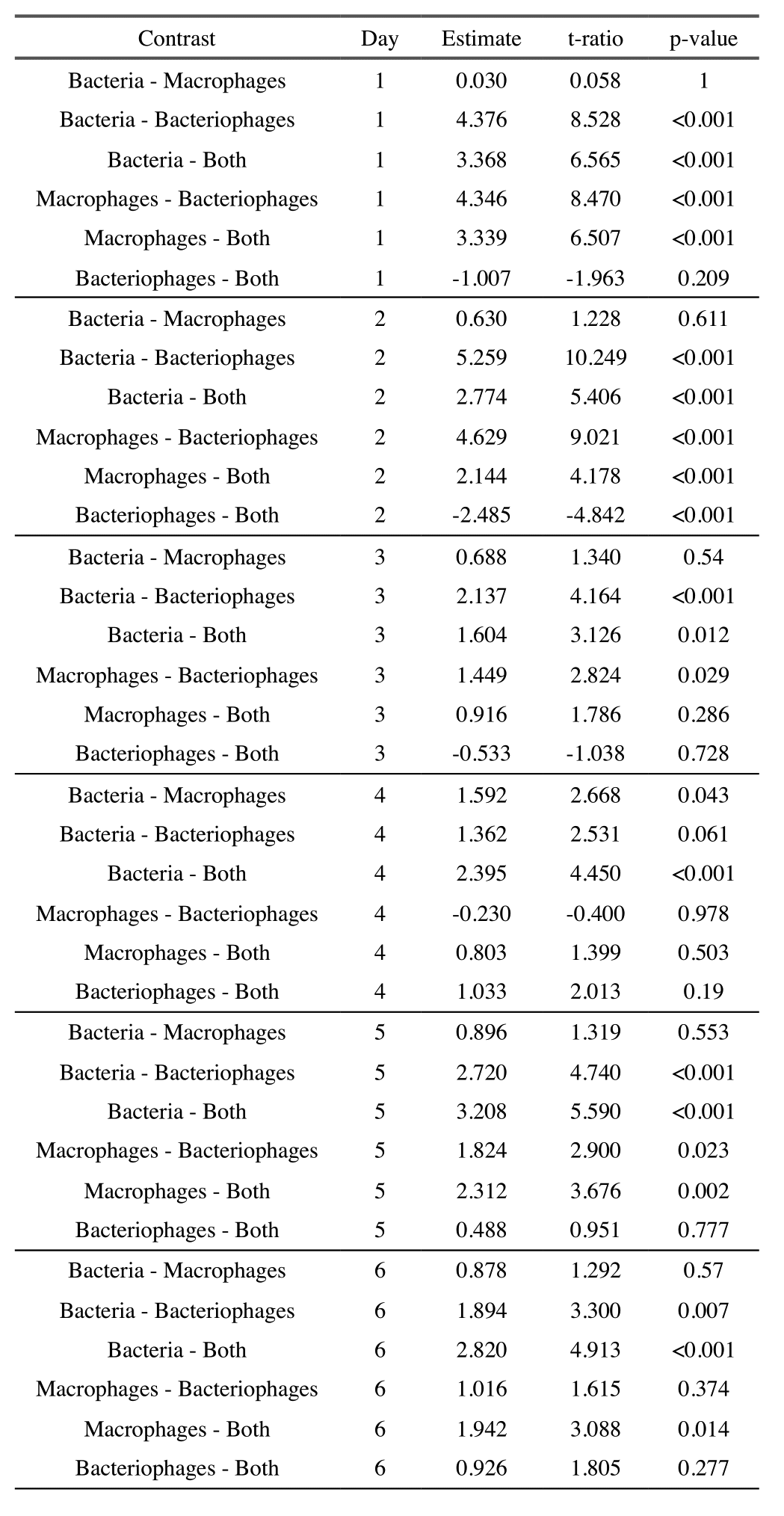

Table S4. Multiple pairwise comparisons comparing bacteriophage density through time (days 1-6). P-values adjusted using the Tukey method of comparing a family of six estimates.

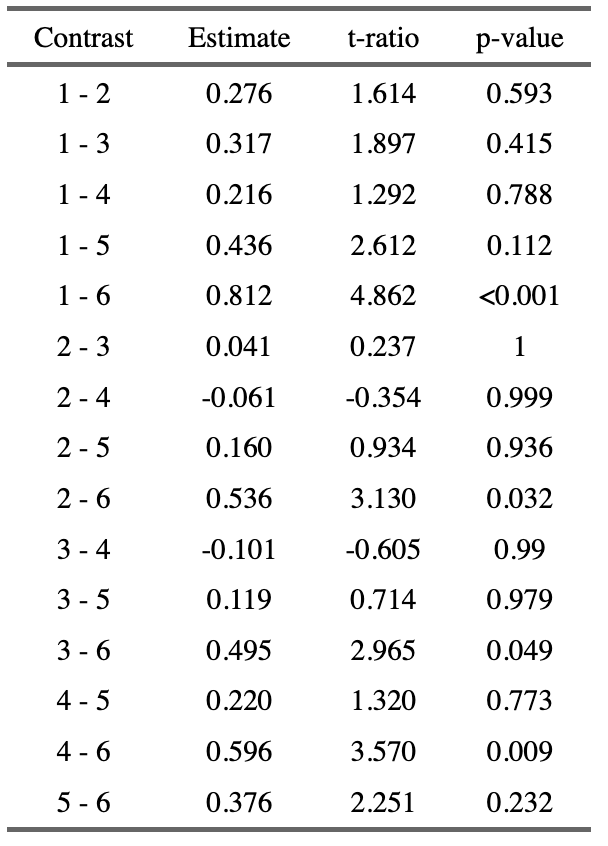

Table S5. Multiple pairwise comparisons comparing bacteriophage growth rates (averaged over 14-1 and PNM as non-significant in the model) in treatments in which the environment was manipulated. The treatment environment included a ‘no macrophage control’ (fresh growth media, no macrophages), ‘macrophages’ (fresh media and unstimulated macrophages), ‘activated media’ (media from macrophages stimulated by heat-killed bacteria) and ‘spent media’ (media isolated from unstimulated macrophage growth). P-values adjusted using the Tukey method of comparing a family of four estimates.

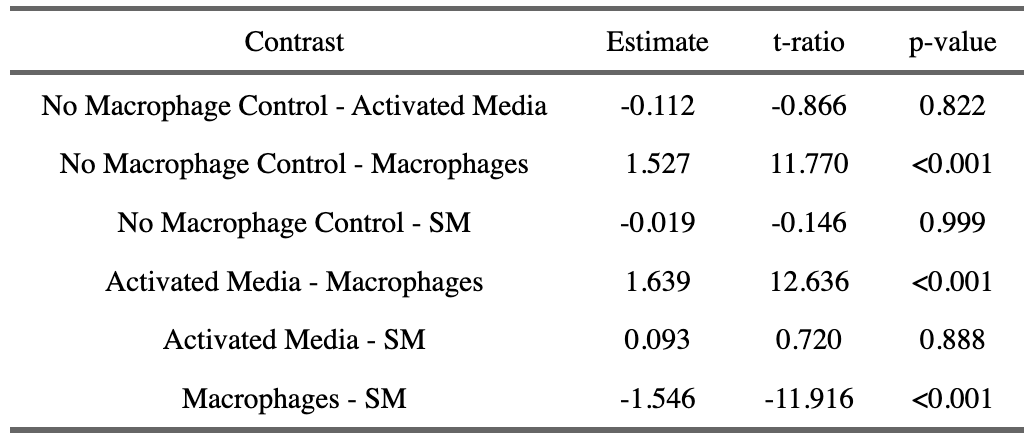

Table S6. Multiple pairwise comparisons comparing biofilm production of bacteria from different evolutionary backgrounds. P-values adjusted using the Tukey method of comparing a family of five estimates.

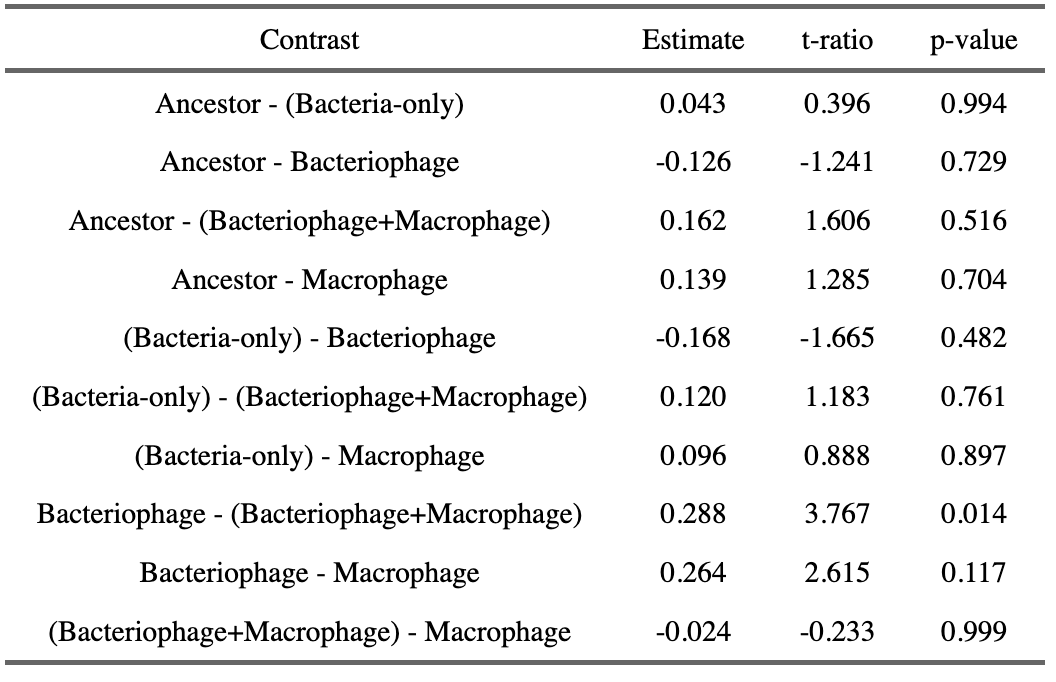

Table S7. Multiple pairwise comparisons comparing cytokine production (TNF-α) of macrophages exposed to bacteria from different evolutionary backgrounds. The macrophage-only control refers to the cytokines produced by macrophages not exposed to bacteria. P-values adjusted using the Tukey method of comparing a family of six estimates.

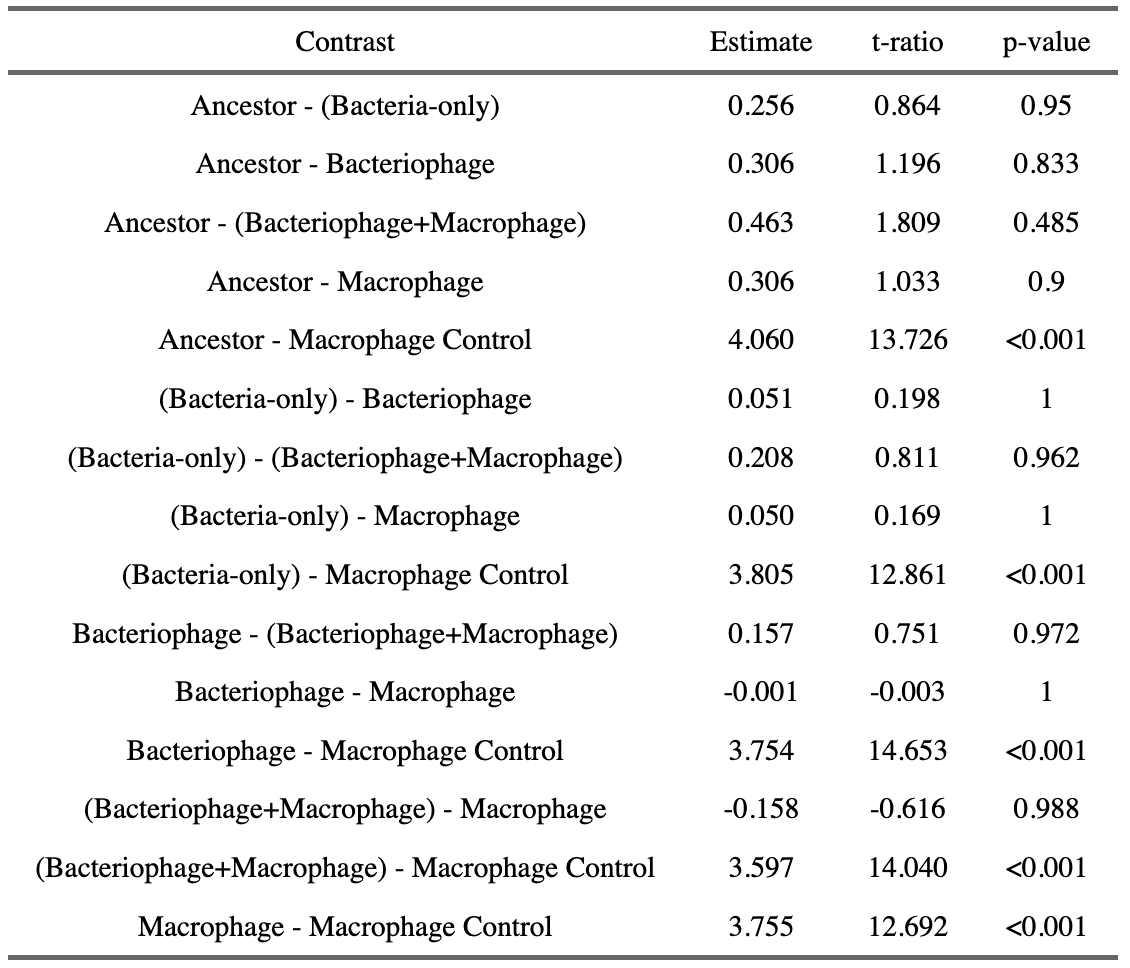
