## Supplementary material for "Antagonism between bacteriophages and macrophages decreases efficacy of a bacteriophage cocktail and increases bacteriophage resistance": Figure S

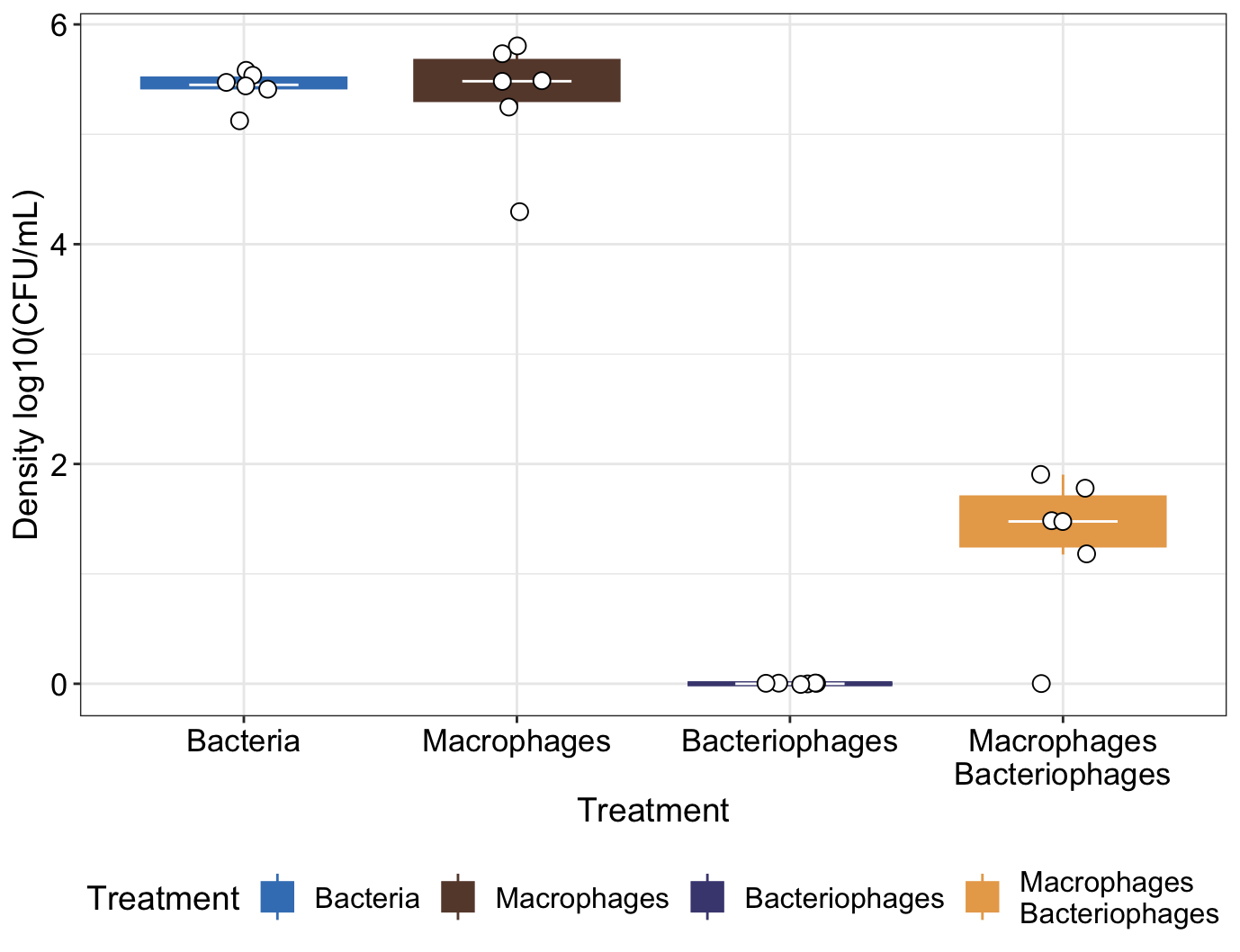
Figure S1. The effects of macrophages and bacteriophages on bacterial density over 8 hrs of bacterial growth. Tops and bottoms of the bars represent the 75th and 25th percentiles of the data, the middle lines are the medians, and the whiskers extend from their respective hinge to the smallest or largest value no further than 1.5* interquartile range.

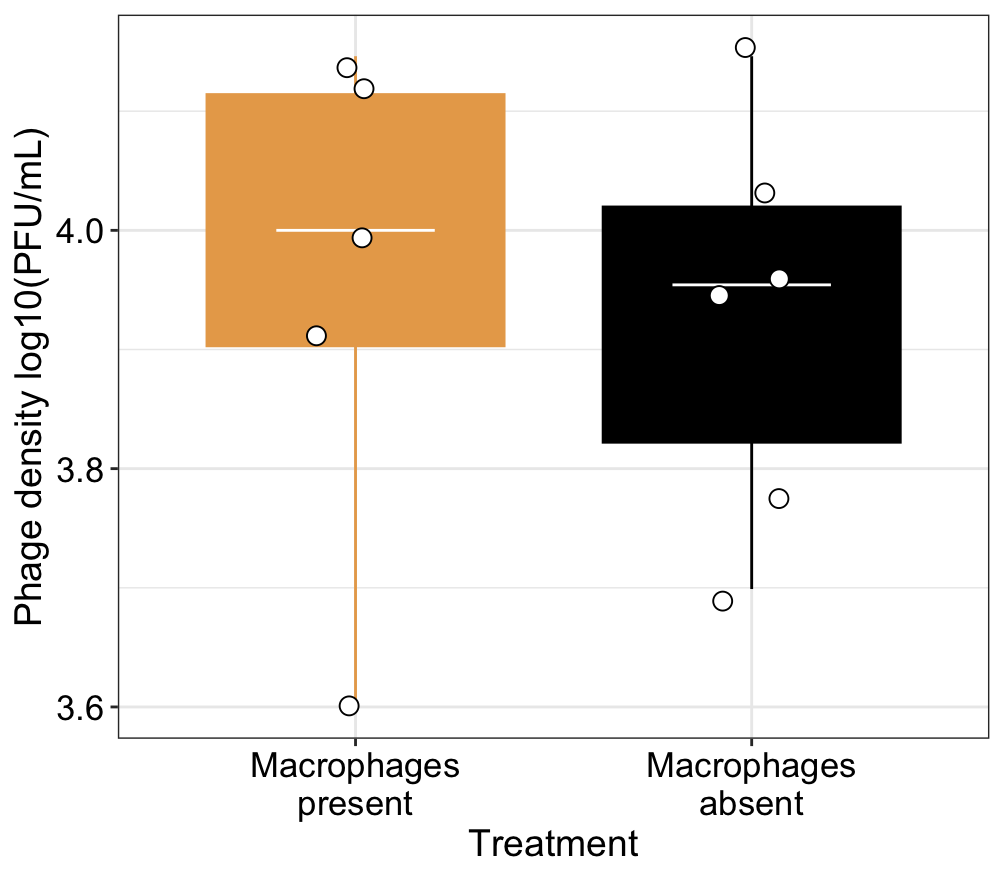

Figure S2. Phage density (log10(PFU/mL)) following culturing with and about macrophages present. Tops and bottoms of the bars represent the 75th and 25th percentiles of the data, the middle lines are the medians, and the whiskers extend from their respective hinge to the smallest or largest value no further than 1.5* interquartile range.

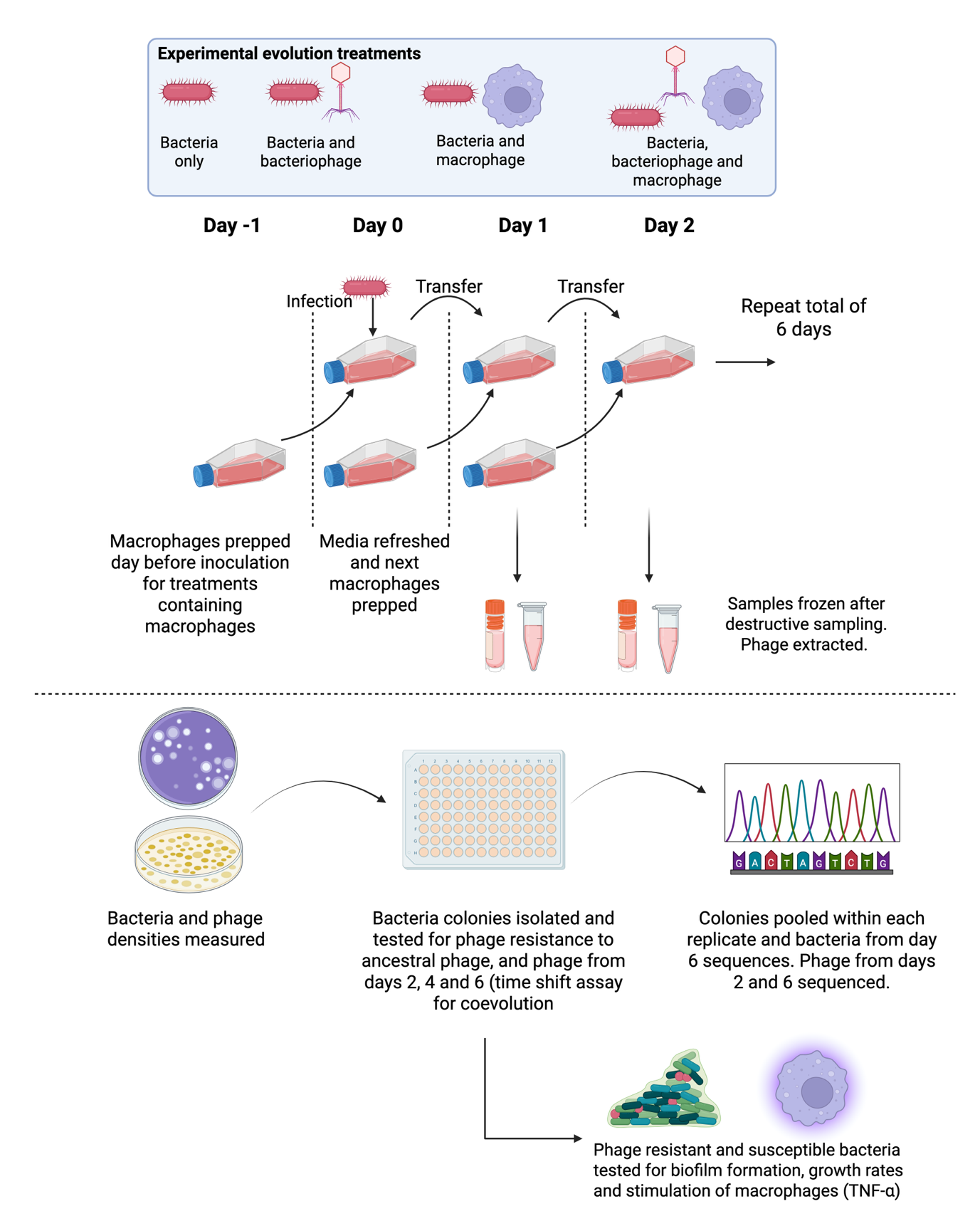

Figure S3. Overview of experimental design for experimental evolution with bacteria, bacteriophages and macrophages. For treatments containing macrophages, macrophages were seeded the day before inoculation/transfer. Bacteria and phage samples were taken daily to measure changes in bacteria and phage density. After experimental evolution, cultures were plated to isolate colonies which were tested for phage resistance in a time-shift assay for measuring coevolution. Isolated colonies were also used for sequencing, biofilm, growth rate and macrophage stimulation analysis.

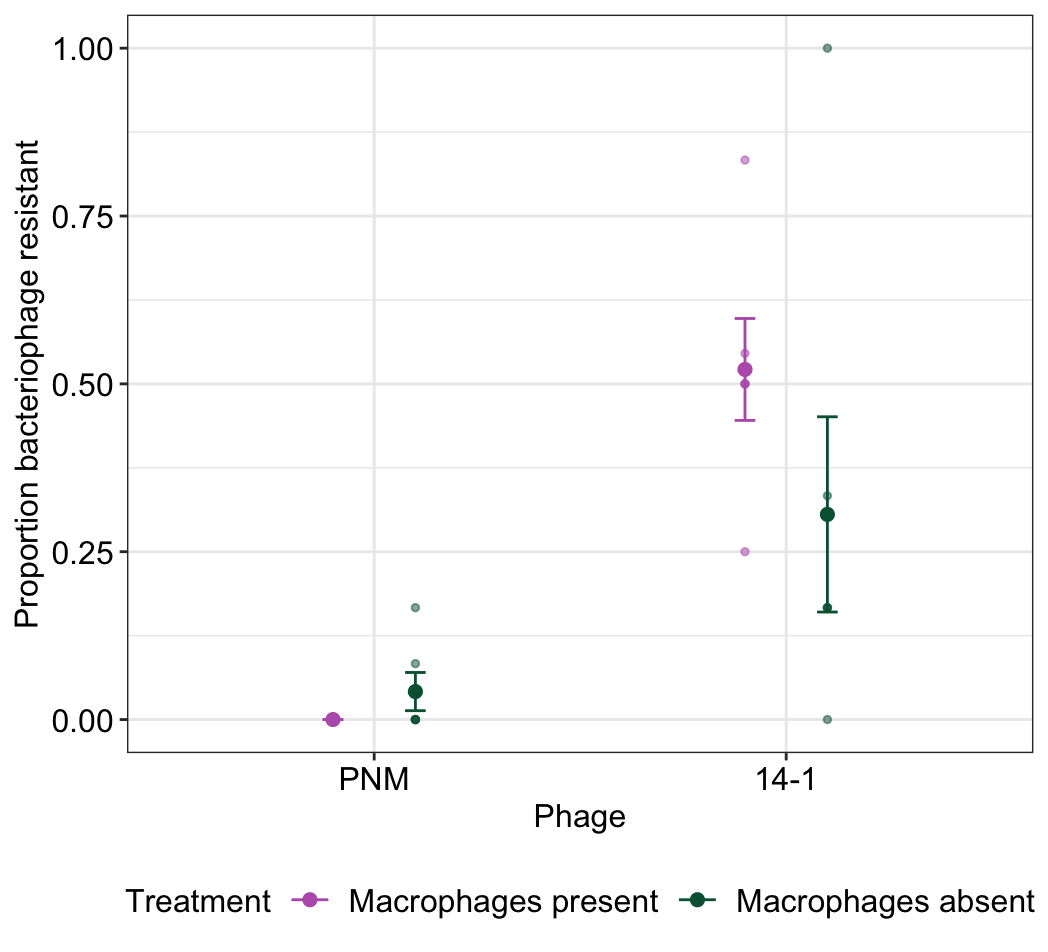

Figure S4. The proportion of isolates within each treatment replicate that were resistant to bacteriophages PNM and 14-1 at day 6 of experimental evolution. Small points indicate individual treatment replicates while larger points with bars indicate means with standard error.

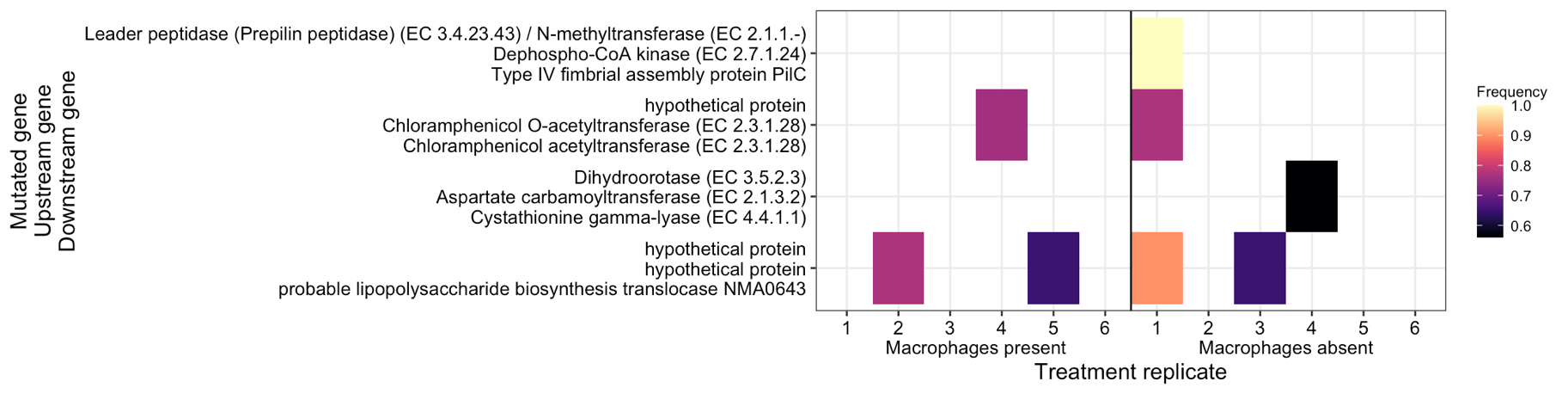

Figure S5. Mutations identified in bacteria populations evolved in the presence of bacteriophages and the presence and absence of macrophages. The mutated gene is presented on the x-axis alongside the upstream and downstream genetic region. Tiles indicate the frequency of the mutation within the population.

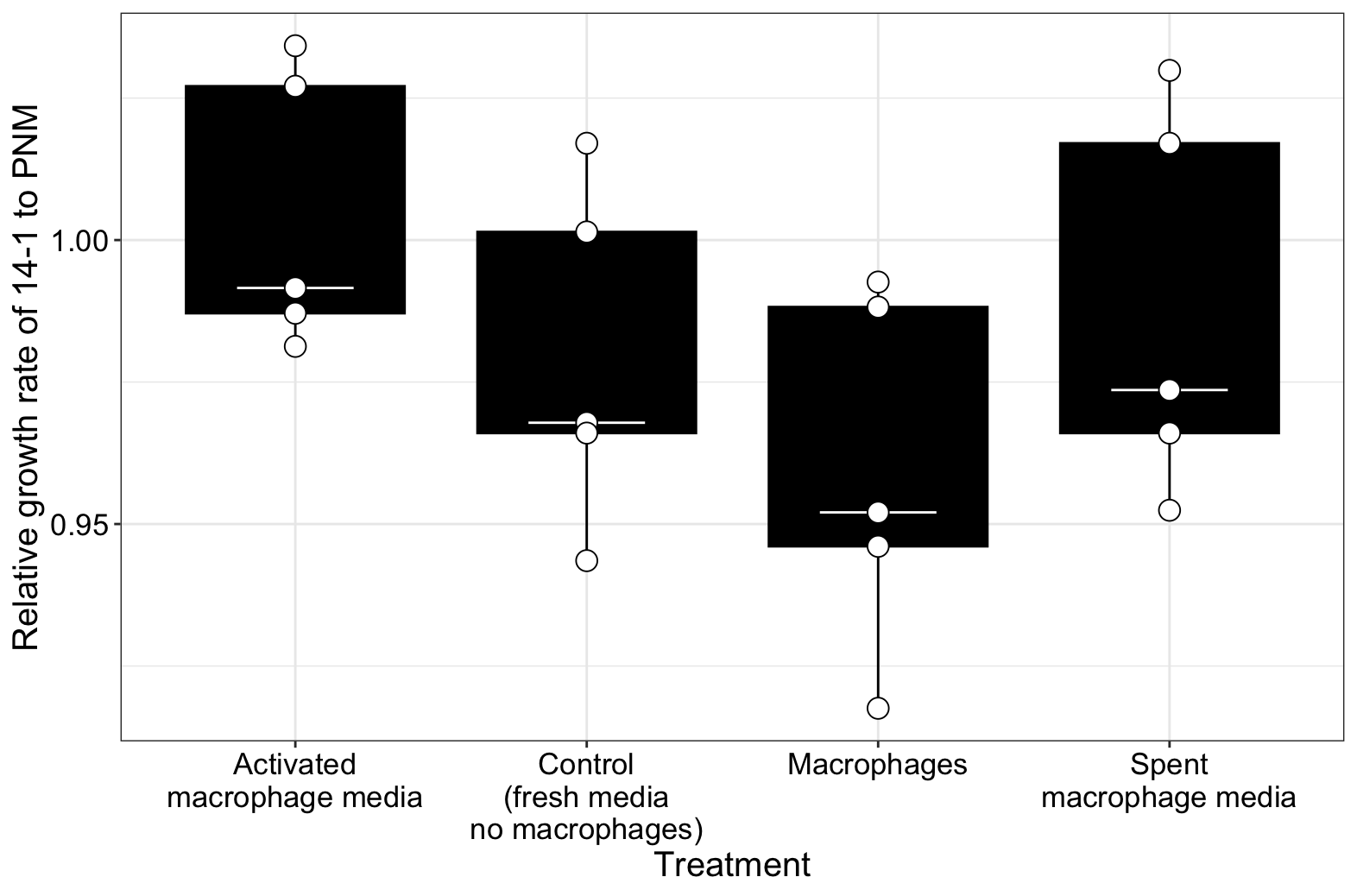

Figure S6. The relative fitness of 14-1 to PNM in different media and macrophage treatments. Points indicate individual treatment replicates. Tops and bottoms of the bars represent the 75th and 25th percentiles of the data, the middle lines are the medians, and the whiskers extend from their respective hinge to the smallest or largest value no further than 1.5* interquartile range.
